# First Molecular-Level Model of the *Chlamydia trachomatis* Injectisome and Targeting CdsN ATPase Oligomerization for Antivirulence Therapy

**DOI:** 10.64898/2026.05.06.723290

**Authors:** Amisha Panda, Jahnvi Kapoor, Raman Rajagopal, Sanjiv Kumar, Anannya Bandyopadhyay

## Abstract

*Chlamydia trachomatis* is an obligate intracellular pathogen that causes sexually transmitted infections and trachoma. Its persistent forms show reduced antibiotic susceptibility, creating a need for new antivirulence strategies. The *C. trachomatis* injectisome or type III Secretion System (T3SS) injects effector proteins into host cells, yet the molecular structure of the complete apparatus is unknown. Here, we identify and validate all 13 T3SS constituent proteins using TXSSScan and reverse-BLAST analysis, and model and assemble them into the complete T3SS apparatus using an integrated computational pipeline. Template-based modeling revealed a conserved architecture despite low sequence identity (18-46%), and the assemblies were validated stereochemically and energetically against homolog controls. Targeting the oligomerization interface of the CdsN ATPase through structure-based virtual screening of the e-Drug3D and IMPPAT libraries, followed by ADMET filtering, including predicted membrane permeability for intracellular targeting, identified three repurposable FDA-approved candidates: M Roflumilast, Elacestrant, and Tecovirimat. MM-GBSA calculations and molecular dynamics simulations confirmed stable complexes with two adjacent CdsN monomers. Elacestrant showed the most favorable binding free energy. These results provide the first molecular-level map of the *C. trachomatis* T3SS and a rational basis for antivirulence drug repurposing.

**Graphical Abstract:** 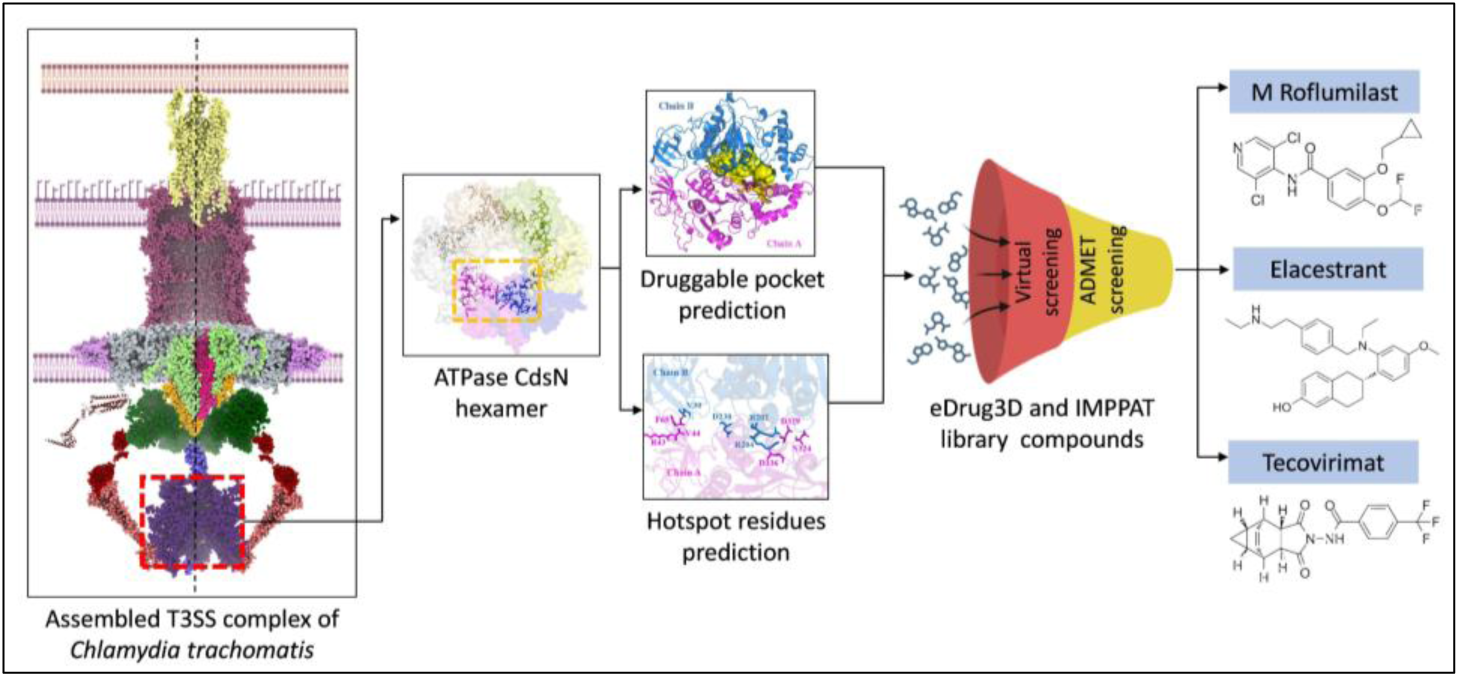

## 1. Introduction

*Chlamydia trachomatis* (Ct) is the leading cause of bacterial sexually transmitted infections (STIs), affecting ∼131 million individuals worldwide annually, with the highest prevalence among individuals aged ≤ 24 years [1]. In the United States alone, the direct lifetime medical costs associated with Ct infections were estimated at approximately $691 million in 2018 [2]. Ct is a Gram-negative, obligate intracellular, non-motile, and ovoid bacterium [3,4]. Although it is transmitted primarily through sexual contact, it can also pass from an infected mother to her newborn during passage through the vaginal canal [5]. Neonates infected in this way may develop trachoma, the foremost cause of infectious blindness globally [6]. Infections are frequently asymptomatic and therefore go undiagnosed; however, persistent or recurrent infections can result in severe complications, including cervicitis, urethritis, pelvic inflammatory disease (PID), cervical cancer, and infertility [7]. The bacterium exhibits a biphasic lifecycle, alternating between the infectious, extracellular elementary body (EB) and the replicative, intracellular reticulate body (RB), and can persist in a metabolically altered state during chronic infection [8]. Because persistent chlamydial forms tolerate multiple antibiotics as a result of their altered physiological states [9], there is an urgent need for alternative antimicrobial strategies against Ct.

An attractive route to such strategies is the injectisome or type III secretion system (T3SS), a highly conserved virulence apparatus employed by many Gram-negative bacteria to drive pathogenesis [10–15]. These pathogens span diverse lifestyles: *Salmonella enterica* and *Shigella flexneri* are facultative intracellular pathogens, whereas *Vibrio cholerae* and *Escherichia coli* are predominantly extracellular. Ct, by contrast, is an obligate intracellular pathogen in which the T3SS is a central virulence determinant, as impairing its activity markedly disrupts intracellular replication and development [16,17]. The system forms a syringe-like proteinaceous channel that spans the bacterial inner (IM) and outer membrane (OM) and projects into the host cell, delivering effector proteins such as TarP (translocated actin recruiting phosphoprotein), TmeA and TmeB (translocated membrane-associated effectors A and B), and Incs (inclusion membrane proteins) [18–20]. Unlike other Gram-negative bacteria, the chlamydial T3SS genes are dispersed across four separate genomic clusters, termed the pathogenicity archipelago, rather than a single contiguous operon within a chromosomal pathogenicity island [21]. This differs from the clustered organization found in other bacteria-the *ipa*, *mxi* and *spa* operons of the *S. flexneri* virulence plasmid [22], the SP-1 and SP-2 pathogenicity islands of *S. enterica* [23], a horizontally acquired genomic island in *V. cholerae* [24], and the locus of enterocyte effacement (LEE) in *E. coli* [25]. Although the overall Ct T3SS architecture is highly conserved with that of other Gram-negative bacteria [26], the apparatus exhibits structural adaptations consistent with its obligate intracellular lifestyle. Ct is thought to have first evolved the T3SS for basic survival and subsequently repurposed it for pathogenesis [26]; accordingly, its needle protein CdsF uniquely harbours cysteine residues that enable the needle to adopt distinct conformations across developmental stages [26]. In situ cryoelectron tomography indicates that host-membrane contact induces basal-body compaction, consistent with conformational rearrangement of the injectisome upon host engagement [27]. The apparatus comprises 13 proteins organized into five subcomplexes-the needle complex, basal body, export apparatus, ATPase complex, and cytoplasmic sorting platform [28] and constitutes a 131-subunit assembly that depends on extensive protein-protein interactions (PPIs). Throughout, we use the unified Sct nomenclature together with the *Chlamydia*-specific Cds designations where appropriate [29]. The needle protein (SctF) forms a central hollow channel for effector translocation [30] and is anchored by the basal body, comprising the outer and inner membrane rings (OMR and IMR) [31]. Embedded within the IM, the export apparatus (SctR, SctS, SctT, SctU, and SctV) functions as an export gate that selects substrates for secretion [32–36]; the ATPase complex (SctN, SctL, and SctO) supplies the energy to unfold and translocate them [28]; and the cytoplasmic sorting platform (SctQ) forms a pod-like structure at the terminus of each SctL dimer [37–40]. Despite this conservation, only a few Ct T3SS proteins have been resolved experimentally-CdsD, CdsO and CdsU [41,42]. Notably, the periplasmic region of CdsD contains an extended α-helical segment absent in other Gram-negative bacteria that is proposed to support basal body assembly and stability [41]. Upon host engagement, the translocators CopB and CopD/CopD2 are secreted and insert into the host membrane to form a translocation pore for effector delivery [43,44].

Because inhibition of T3SS function attenuates chlamydial replication and development [16,17], its constituent proteins are attractive targets for anti-chlamydial agents. The export apparatus is powered by the ATPase CdsN, which harnesses cooperative ATP hydrolysis at inter-subunit catalytic sites to unfold and extract substrates [26,45,46]. In an earlier study, Grishin et al. screened a homology model of Ct CdsN at its catalytic site and identified two compounds that impaired chlamydial development and suppressed T3SS-mediated secretion of the effector IncA [47]. Targeting inter-subunit contacts, particularly the monomer-monomer interface, represents a complementary strategy for inhibiting ATPase activity and attenuating virulence [48].

Although the individual Ct T3SS proteins have been catalogued, the architecture of the complete Ct T3SS remains undefined. Indeed, no complete T3SS structure has yet been fully resolved in any Gram-negative bacterium, as the experimental characterization of such large multiprotein assemblies remains technically challenging. Computational modelling therefore provides a valuable complementary approach [49]. Here, we compiled all 13 Ct T3SS proteins from the literature and retrieved their sequences using the corresponding locus tags. Sequences were validated as T3SS components via reverse BLAST and TXSS Scan tool. Monomeric models were obtained from the AlphaFold database and, guided by structural homology with the T3SS architectures of other Gram-negative bacteria, assembled into oligomeric subcomplexes, for which binding affinities and intermolecular interactions were determined. Rather than the conserved catalytic motifs targeted previously in Ct and other pathogens [47,50,51], we focused on the Ct CdsN oligomerization interface, screening compounds from the e-Drug3D and IMPPAT libraries against its predicted hotspots and druggable pocket, filtering the hits by pharmacokinetic (ADMET) criteria, and evaluating binding energetics and complex stability using MM-GBSA and molecular dynamics (MD) simulations. Together, these analyses provide the first molecular-level model of the Ct T3SS and identify three candidate compounds with the potential to attenuate Ct virulence.

## 2. Materials and Methods

### 2.1 Retrieval of T3SS constituent proteins and sequence alignment with corresponding homologs

We selected the reference strain Ct D/UW-3/CX for this study. 13 T3SS constituent proteins were obtained through literature survey [26]. The computational framework used in this study builds upon and extends the methodology described by Kapoor et al. (2026) [49]. A detailed description of the methodology used in this study is provided below, and the overall workflow is schematically represented in Figure 1. The FASTA sequences of 13 T3SS proteins were retrieved from UniProt using their gene locus tags (gene identifiers). We further validated the identified proteins as T3SS constituents using TXSSScan tool [52] (https://galaxy.pasteur.fr/) (accessed on 3 September 2025) and a reverse BLAST approach against the Ct D/UW-3/CX proteome.

**Figure 1:**
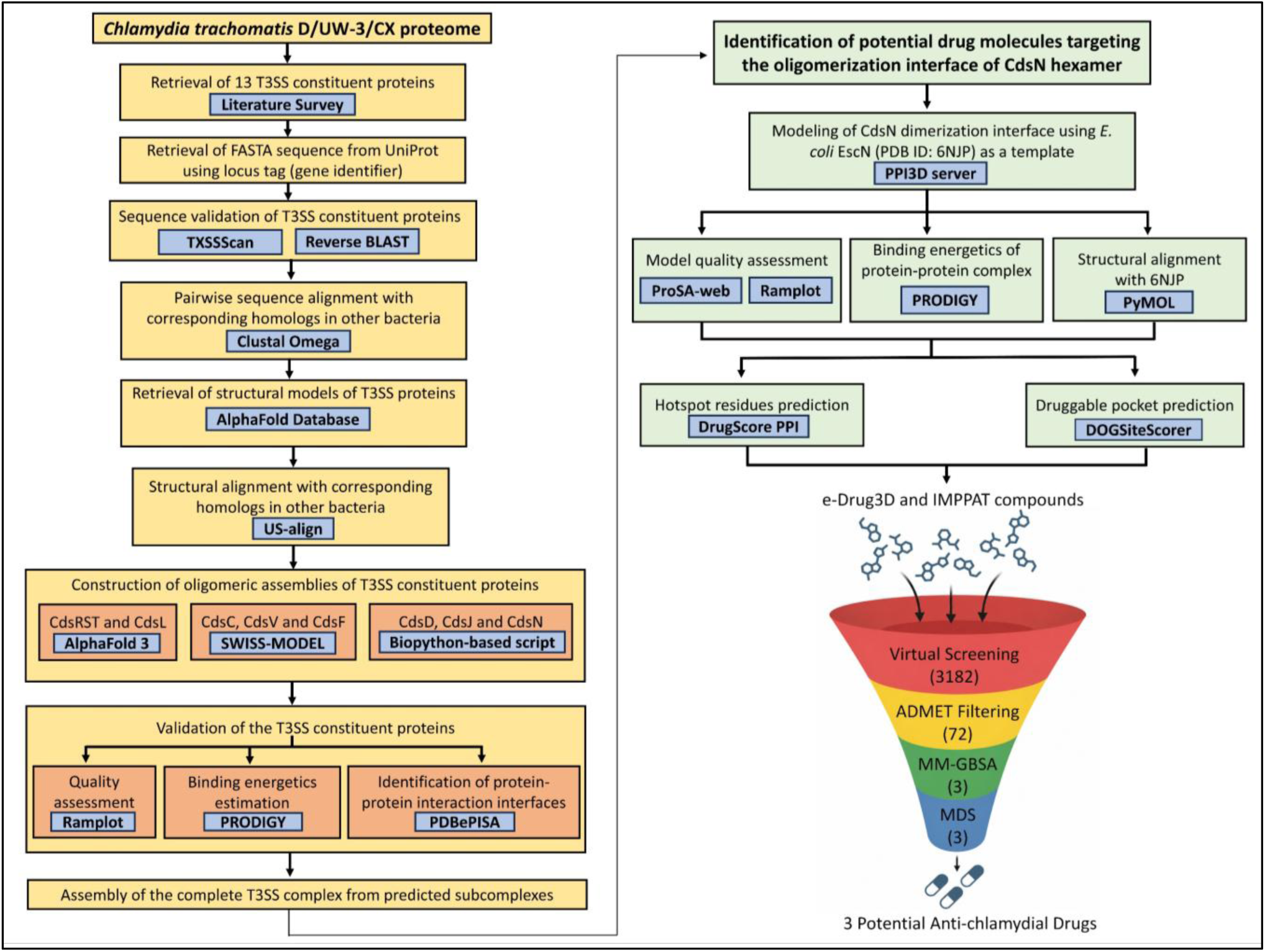
Computational framework for the structural characterization and assembly of T3SS constituent proteins in Ct D/UW-3/CX proteome, followed by structure-based virtual screening of compounds targeting the oligomerization interface of CdsN hexamer. Virtual screening, ADMET filtering, MM-GBSA analysis, and molecular dynamics simulations were performed using the Schrodinger Suite 2025-3.

Following validation, pairwise sequence alignment was performed with their respective T3SS homologs in other Gram-negative bacteria (*S. enterica*, *S. flexneri*, *E. coli* and *V. cholerae*) using Clustal Omega [53] (https://www.ebi.ac.uk/jdispatcher/msa/clustalo) (accessed on 5 September 2025) to assess the degree of sequence similarity (Table 1).

**Table 1:**
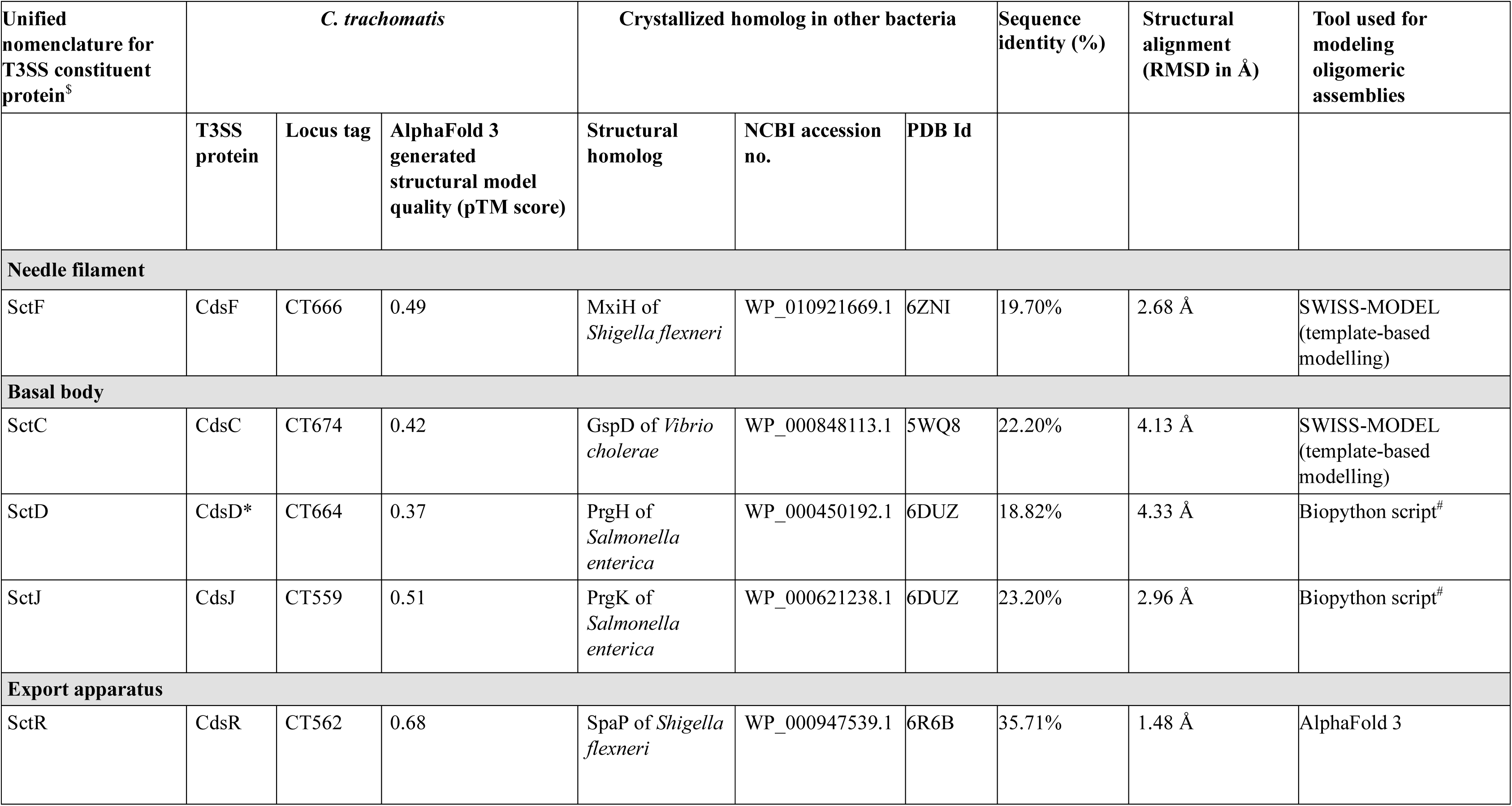
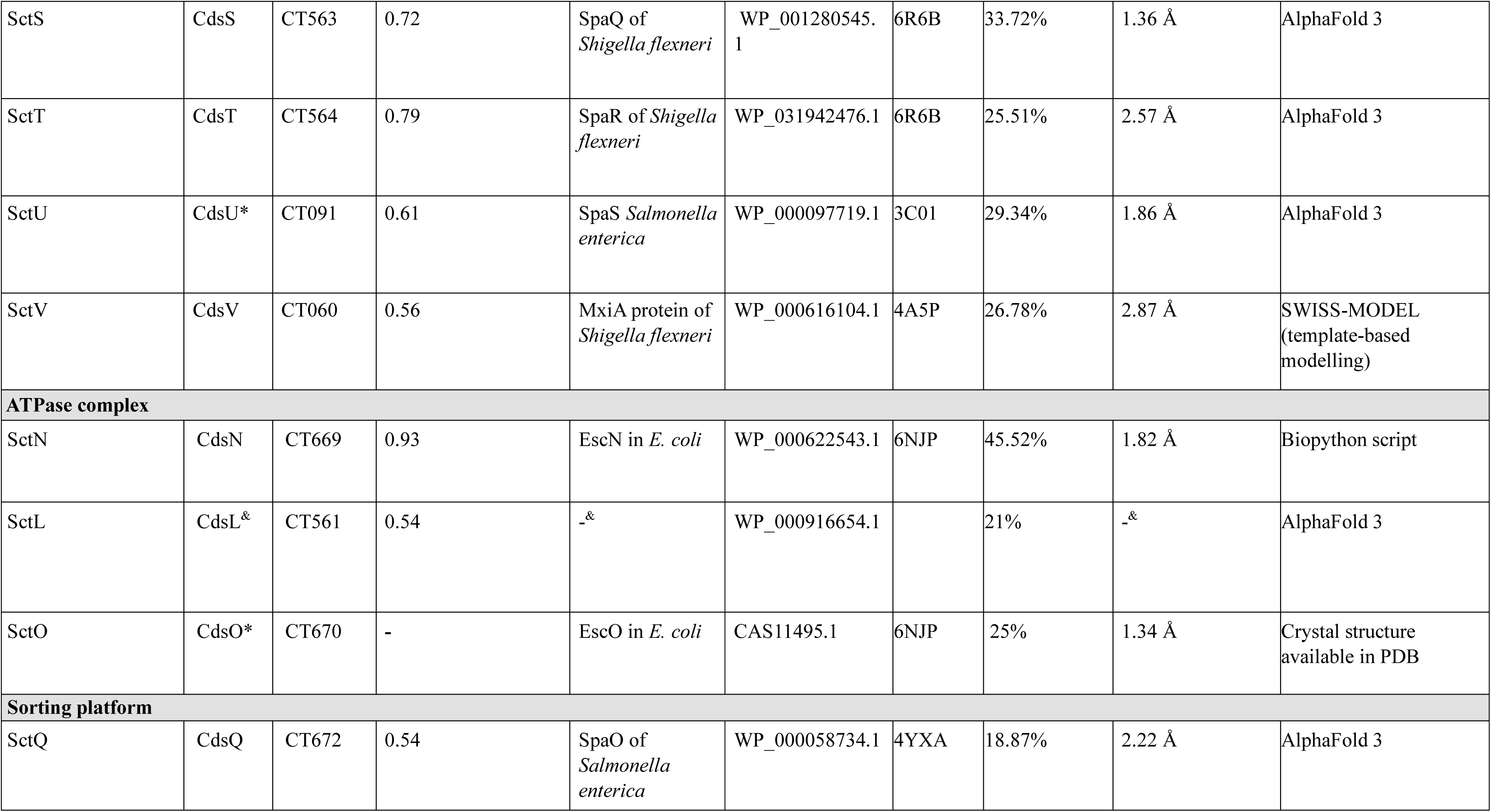
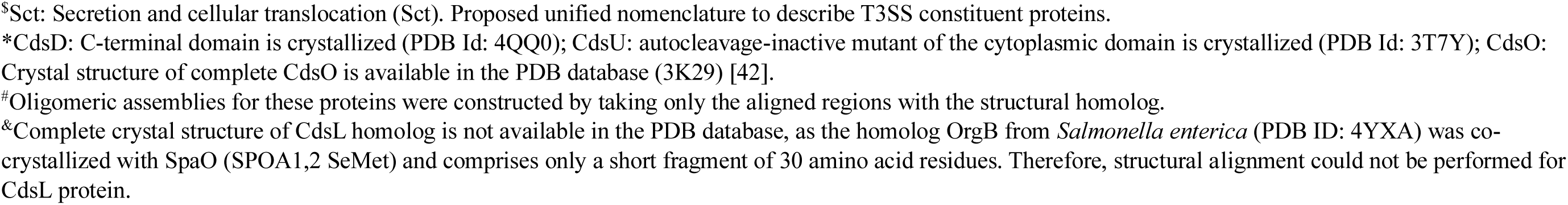
Sequence and structural similarity of *C. trachomatis* T3SS constituent proteins with their homologs in other Gram-negative bacteria.

| Unified nomenclature for T3SS constituent protein<sup>s</sup> | *C. trachomatis* |  |  | Crystallized homolog in other bacteria |  |  | Sequence identity (%) | Structural alignment (RMSD in Å) | Tool used for modeling oligomeric assemblies |
| --- | --- | --- | --- | --- | --- | --- | --- | --- | --- |
|  | T3SS protein | Locus tag | AlphaFold 3 generated structural model quality (pTM score) | Structural homolog | NCBI accession no. | PDB Id |  |  |  |
| **Needle filament** |  |  |  |  |  |  |  |  |  |
| SctF | CdsF | CT666 | 0.49 | MxiH of *Shigella flexneri* | WP_010921669.1 | 6ZNI | 19.70% | 2.68 Å | SWISS-MODEL (template-based modelling) |
| **Basal body** |  |  |  |  |  |  |  |  |  |
| SctC | CdsC | CT674 | 0.42 | GspD of *Vibrio cholerae* | WP_000848113.1 | 5WQ8 | 22.20% | 4.13 Å | SWISS-MODEL (template-based modelling) |
| SctD | CdsD* | CT664 | 0.37 | PrgH of *Salmonella enterica* | WP_000450192.1 | 6DUZ | 18.82% | 4.33 Å | Biopython script<sup>#</sup> |
| SctJ | CdsJ | CT559 | 0.51 | PrgK of *Salmonella enterica* | WP_000621238.1 | 6DUZ | 23.20% | 2.96 Å | Biopython script<sup>#</sup> |
| **Export apparatus** |  |  |  |  |  |  |  |  |  |
| SctR | CdsR | CT562 | 0.68 | SpaP of *Shigella flexneri* | WP_000947539.1 | 6R6B | 35.71% | 1.48 Å | AlphaFold 3 |
| SctS | CdsS | CT563 | 0.72 | SpaQ of *Shigella flexneri* | WP_001280545.1 | 6R6B | 33.72% | 1.36 Å | AlphaFold 3 |
| SctT | CdsT | CT564 | 0.79 | SpaR of *Shigella flexneri* | WP_031942476.1 | 6R6B | 25.51% | 2.57 Å | AlphaFold 3 |
| SctU | CdsU* | CT091 | 0.61 | SpaS *Salmonella enterica* | WP_000097719.1 | 3C01 | 29.34% | 1.86 Å | AlphaFold 3 |
| SctV | CdsV | CT060 | 0.56 | MxiA protein of *Shigella flexneri* | WP_000616104.1 | 4A5P | 26.78% | 2.87 Å | SWISS-MODEL (template-based modelling) |
| **ATPase complex** |  |  |  |  |  |  |  |  |  |
| SctN | CdsN | CT669 | 0.93 | EscN in *E. coli* | WP_000622543.1 | 6NJP | 45.52% | 1.82 Å | Biopython script |
| SctL | CdsL<sup>&</sup> | CT561 | 0.54 | -<sup>&</sup> | WP_000916654.1 |  | 21% | -<sup>&</sup> | AlphaFold 3 |
| SctO | CdsO* | CT670 | - | EscO in *E. coli* | CAS11495.1 | 6NJP | 25% | 1.34 Å | Crystal structure available in PDB |
| **Sorting platform** |  |  |  |  |  |  |  |  |  |
| SctQ | CdsQ | CT672 | 0.54 | SpaO of *Salmonella enterica* | WP_000058734.1 | 4YXA | 18.87% | 2.22 Å | AlphaFold 3 |

<sup>\$</sup>Sct: Secretion and cellular translocation (Sct). Proposed unified nomenclature to describe T3SS constituent proteins.
\*CdsD: C-terminal domain is crystallized (PDB Id: 4QQ0); CdsU: autocleavage-inactive mutant of the cytoplasmic domain is crystallized (PDB Id: 3T7Y); CdsO: Crystal structure of complete CdsO is available in the PDB database (3K29) [42].

### 2.2 Construction of structural models of T3SS constituent proteins

Monomeric structural models of the T3SS proteins were retrieved from the AlphaFold database (https://alphafold.ebi.ac.uk/) (accessed on 7 September 2025) [54]. The quality and reliability of the predicted structural models were assessed using AlphaFold 3 confidence metrics, including the pLDDT (predicted Local Distance Difference Test), PAE (Predicted Aligned Error), pTM (predicted Template Modeling), and ipTM (inter-chain predicted TM) scores. For cross-validation, structural models of proteins having pTM scores below 0.5 (CdsF, CdsC, and CdsD) were further generated using four additional tools-ESMFold [55], SWISS-MODEL [56], TrRosetta [57] and RoseTTAFold2 [58]. The resulting atomic coordinate files were visualized in PyMOL 3.0 [59]. AlphaFold 3-predicted structural models of monomeric proteins were used for further analysis.

### 2.3 Structural comparison of T3SS proteins with crystal structures of their homologs

We downloaded the atomic coordinate files of the corresponding T3SS homologs in other Gram-negative bacteria from the RCSB PDB database (https://www.rcsb.org/) (accessed on 9 September 2025) with PDB Ids mentioned in Table 1. Structural alignment was performed between the Ct T3SS proteins and corresponding homologs using the US-align online server [60] (https://aideepmed.com/US-align/) (accessed on 9 September 2025). The resulting RMSD (Root Mean Square Deviation) values were used to validate the predicted structural models (Table 1).

### 2.4 Construction of oligomeric assemblies

The complete architecture of the Ct T3SS can be organized into five major subcomplexes, arranged from outside to inside across the bacterial cell membrane: the needle complex, basal body, export apparatus, ATPase complex, and the cytoplasmic sorting platform. Oligomeric assemblies of each T3SS constituent protein were constructed using three complementary approaches: AlphaFold 3, template-based modeling with SWISS-MODEL, and a Biopython-based script. Using AlphaFold 3 (accessed on 10 September 2025), structural models of CdsL dimer and the CdsRST complex were generated [61]; however, this approach could not be applied to other T3SS oligomeric proteins due to sequence length limitations in AlphaFold 3 server [62]. Therefore, template-based modeling using SWISS-MODEL (https://swissmodel.expasy.org/) (accessed on 13 September 2025) [56] was used to generate oligomeric structures of CdsF, CdsC and CdsV based on the corresponding structural templates listed in Table 1. As SWISS-MODEL could not predict oligomeric assemblies for the remaining T3SS constituent proteins, a Biopython-based script [49,63] was used to assemble oligomeric structural models of CdsD-CdsJ and CdsN. This script aligned and assembled multiple protein subunits into a single complex using crystallographic structures of homologous proteins from other Gram-negative bacteria as templates (Table 1) to define spatial arrangement and orientation.

All modeled subcomplexes were visualized in UCSF ChimeraX 1.9 using corresponding atomic coordinate files of the T3SS proteins [64]. The subcomplexes were manually assembled based on the predicted Ct T3SS model and structural organization of T3SS architecture in *S. enterica* and *S. flexneri* to constitute the overall architecture of Ct T3SS [14,26].

### 2.5 Quality assessment of oligomeric assemblies

The overall quality of each generated structural model of T3SS proteins was assessed using Ramplot (https://www.ramplot.in/) (accessed on 20 September 2025) [65]. Ramplot is an online tool that generates both standard 2D and advanced 3D Ramachandran plots, which enables visualization of allowed and disallowed conformational regions for a given protein or peptide [65]. The structural models of subcomplexes were further validated by calculating binding energetics (ΔG, kcal/mol) and dissociation constant (K_d_, M) using the PRODIGY (PROtein binDIng enerGY prediction) server (https://rascar.science.uu.nl/prodigy/) (accessed on 20 September 2025) [66], which predicts the binding affinity of protein-protein complexes. ΔG and K_d_ values were also calculated for crystallographic structures of homologous T3SS proteins, which served as positive controls for validation of predicted oligomeric assemblies. In addition, PDBePISA (https://www.ebi.ac.uk/msd-srv/prot_int/cgi-bin/piserver) (accessed on 22 September 2025) [67] was used to predict the amino acid residues involved in protein-protein interactions, including hydrogen bonds, salt bridges, and hydrophobic contacts. PDBePISA (Proteins, Interfaces, Structures and Assemblies) is an online tool for analyzing macromolecular interfaces, which provides insights into the structural and chemical properties of interacting surfaces.

### 2.6 Homology modeling of CdsN ATPase

Because CdsN functions as a homohexamer in the assembled T3SS, but our virtual screening targets the inter-subunit PPI interface, two adjacent monomers were used to represent the oligomerization interface; the term ‘dimer’ is used hereafter to denote this pair of adjacent CdsN monomers. We submitted the CdsN sequence to the AlphaFold 3 server as a dimeric query; however, the predicted model exhibited low confidence, as reflected by low ipTM score (0.16). Therefore, the dimeric structure of Ct CdsN was generated through homology modeling using the PPI3D (Protein-Protein Interactions in Three Dimensions) server (https://bioinformatics.lt/ppi3d/) (accessed on 24 September 2025) [68], which integrates MODELLER software for structure prediction. PPI3D enables users to identify interactions between a pair of proteins and perform template-based homology modeling by leveraging homologous structural information from available databases [68]. We submitted the amino acid sequence of CdsN to the PPI3D as a dimeric query, and the option to search for close homologs using BLAST was enabled. The ATPase EscN from *E. coli* (PDB ID: 6NJP) was identified as the closest homolog, with 47% sequence identity and 3% gap coverage. Based on this template, PPI3D constructed a three-dimensional model of the CdsN dimer, which was visualized using PyMOL software (accessed on 25 September 2025) [59]. The generated CdsN dimer structural model was superimposed onto the crystal structure of the *E. coli* EscN dimer in PyMOL (accessed on 27 September 2025) [69], and the corresponding RMSD value was calculated to assess the structural similarity.

Furthermore, in order to assess the oligomerization interface of CdsN hexamer as a drug target, CdsN homology was checked with the human proteome (TaxID: 9606) using BLASTP (E-value < 1.0E-03, bitscore > 100) (https://blast.ncbi.nlm.nih.gov/Blast.cgi?PROGRAM=blastp&PAGE_TYPE=BlastSearch&LINK_LOC=blasthome) (accessed on 28 September 2025).

### 2.7 Quality assessment of CdsN ATPase

The quality of the generated CdsN dimeric structural model was assessed using the ProSA-web server (https://prosa.services.came.sbg.ac.at/prosa.php) (accessed on 1 October 2025) [70] and Ramplot (https://www.ramplot.in/) (accessed on 3 October 2025) [71]. The ProSA (Protein Structure Analysis) web server generates a Z-score for the input protein model based on its comparison with the known experimentally determined structures in the Protein Data Bank (PDB) [70]. The model was validated based on Z-score and the distribution of amino acid residues in the Ramachandran plot, which provide insights into structural stability and stereochemical quality of the predicted structure, respectively.

The binding energetics of CdsN were evaluated using the PRODIGY webserver (accessed on 25 September 2025) [66]. We submitted the PDB coordinate file of CdsN dimer, generated by the PPI3D server, as input. The server then generated the predicted binding free energy (ΔG) in kcal/mol, dissociation constant (K_d_) in M, along with the number and types of intermolecular contacts within the 5.5 Å distance cutoff, and the percentage of charged and polar amino acids on the non-interacting surface.

### 2.8 Prediction of hotspot residues and druggable pocket

DrugScore PPI (https://cpclab.uni-duesseldorf.de/dsppi/main.php) (accessed on 18 October 2025) [72] was used to identify hotspot residues at the protein-protein interface. It employs a computational approach to predict changes in binding free energy (ΔΔG) resulting from alanine mutations at interface residues. We submitted the PDB coordinate file of the CdsN dimer and specified the chain to be mutated. Hotspot residues were then predicted independently for each chain. Residues showing ΔΔG > 1 kcal/mol were classified as ‘hotspots’, which suggests their essential role in stabilizing the protein-protein interaction.

DOGSiteScorer (https://proteins.plus/help/dogsite) (accessed on 19 October 2025) [73] was used to identify the druggable pocket at the CdsN dimeric interface. The tool accepts a PDB coordinate file as input and predicts potential binding pockets, along with their corresponding druggability scores using a support vector machine (SVM)-based approach. The druggability score ranges from 0 to 1, where higher values indicate a higher likelihood of the pocket being druggable [73].

### 2.9 Molecular docking

The structural model of CdsN dimer was prepared using the Protein Preparation Wizard in Schrodinger Suite 2025-3 [74], followed by energy minimization using the OPLS4 force field. The receptor grids were then generated around the predicted hotspot residues and druggable pocket in the prepared CdsN dimer. The hotspot residues specified for grid generation are shown in Figure 6b, while the residues defining the druggable pocket are shown in Figure 6c.

e-Drug3D (https://chemoinfo.ipmc.cnrs.fr/MOLDB/index.php) (accessed on 26 October 2025) [75] and IMPPAT 2.0 (https://cb.imsc.res.in/imppat/) (accessed on 29 October 2025) [76] databases were used for virtual screening. SD (Structure-Data) files of three-dimensional structures of 2162 drug compounds were downloaded from the e-Drug3D database [75]. SD files of three-dimensional structures of 1020 phytochemicals were downloaded from the IMPPAT database on the basis of drug likeness properties such as Lipinski’s rule of five (Ro5) (zero violation), Ghose rule (zero violation), GSK4/40 (Good), Pfizer: 3/75 (Good), Veber rule (Good), Egan rule (Good) and QEDw (> 0.5). Ligand molecules downloaded from the respective databases were prepared for docking using the LigPrep module of the Schrodinger suite 2025-3.

The prepared ligand molecules were docked onto the predicted interface hotspot regions and druggable pocket of the CdsN dimer using the Glide module [77] of the Schrodinger suite 2025-3. The algorithm identifies favorable hydrogen-bonding, hydrophobic, and electrostatic interactions while penalizing steric hindrance. Following minimization, the docked poses are re-evaluated and ranked using the GlideScore scoring function [77]. Ligand molecules from both databases exhibiting docking scores ≤ −7.0 kcal/mol (Tables S3-S6) were selected for ADMET screening.

### 2.10 ADMET screening

The ADMET properties of the selected ligands from both databases with docking scores ≤ −7.0 kcal/mol were evaluated using the QikProp module of the Schrodinger suite 2025-3, followed by their filtration based on pharmacokinetic properties (Tables S3-S6). Eight criteria were applied for screening: one violation of Lipinski’s Rule of Five, ≥80% predicted human oral absorption, QPPCaco > 500, QPPMDCK (<25 poor, 25-500 moderate and >500 excellent), QPlogHERG < −5, donor HB ≤ 5, acceptor HB ≤ 10, and QPlogPo/w (−2 to 6.5). The QikProp user manual was used to determine acceptable ranges for each property [78]. The three protein-ligand complexes from the e-Drug3D database were selected for evaluation of binding energetics and MD simulation.

### 2.11 Estimation of binding free energy

The binding free energies (ΔG_bind_) of the selected protein-ligand complexes were calculated using the Prime module of the Schrodinger suite 2025-3 through the molecular mechanics-generalized born surface area (MM-GBSA) approach. This method employed the optimized potentials for liquid simulation (OPLS4) force field and VSGB solvent model, along with search algorithms [79]. The binding free energy was determined using the following relationship:

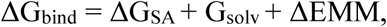

where ΔG_SA_ represents the change in surface area energies between the protein-ligand complexes and the individual protein and ligands, ΔG_solv_ represents the difference in the solvation energies of the complexes and the individual protein and ligands, and ΔEMM reflects the difference in molecular mechanics energies between the protein-ligand complexes and the individual protein and ligands when minimized.

### 2.12 MD simulation

MD simulations were performed to assess the structural stability of the selected protein-ligand complexes using the Desmond module in the Schrodinger suite 2025-3. The docked protein-ligand complexes were first prepared using the System Builder tool, where each protein-ligand complex was solvated with TIP3P water molecules and enclosed with an orthorhombic simulation box. The OPLS4 force field at physiological pH 7.4 was applied, and Na+ and Cl-ions were added to neutralize the system and maintain a salt concentration of 0.15M.

To capture local dynamics and conformational changes associated with ligand binding, we performed short-length simulations. Four independent iterative MD simulation runs of 50 ns each were performed for each protein-ligand complex under NPT conditions at a bar pressure of 1.01325 and a constant temperature of 300K, resulting in a cumulative simulation time of 200 ns per system. In this iterative approach, the final frame from every 50 ns simulation was used as the starting structure for subsequent simulations. As a backup, we also performed an independent 100 ns MD simulation for each protein-ligand complex. The Simulation Interactions Diagram (SID) tool was subsequently used to analyze the trajectories, which generated RMSD and root mean square fluctuation (RMSF) plots, and detailed protein-ligand interaction profiles for each trajectory frame of the simulation.

## 3. Results

### 3.1 Retrieval and homology analysis of T3SS constituent proteins

The 13 T3SS constituent proteins of the Ct D/UW-3/CX reference genome were compiled by literature survey [26], and their FASTA sequences were retrieved from UniProt using the corresponding locus tags (Table 1). For validation, TXSSScan analysis recovered only eight proteins (CdsC, CdsJ, CdsR, CdsS, CdsT, CdsV, CdsN and CdsQ). Because TXSSScan relies on conserved Hidden Markov Model (HMM) profiles, distinct genomic organization and divergent sequences of Ct T3SS likely limit detection of all 13 T3SS constituent proteins [26]. Reverse-BLAST against the reference genome therefore confirmed all 13 proteins (CdsF, CdsC, CdsD, CdsJ, CdsR, CdsS, CdsT, CdsU, CdsV, CdsN, CdsL, CdsO and CdsQ), each at 100% identity.

Pairwise alignment showed that apart from CdsR, CdsS and CdsN, all proteins exhibited < 30% sequence similarity with their homologs (*S. enterica*, *S. flexneri*, *E. coli* and *V. cholerae*), indicating substantial sequence divergence. Monomeric structural models of the proteins retrieved from the AlphaFold database were all reliable, meeting pTM > 0.5 (or pLDDT > 70 over most regions for CdsF, CdsC and CdsD; Table 1) [80]. For the three proteins with low pTM scores (CdsF, CdsC and CdsD), cross-validation with four additional structure prediction tools (Table S1) supported the overall CdsF fold (low RMSD), whereas CdsC and CdsD showed higher RMSD (5.40 Å and 5.82 Å, respectively), attributable to extensive intrinsically disordered regions that different algorithms model divergently (Figure S1). Consistent with the principle that homologous proteins retain a common structural fold over evolutionary time scales despite sequence drift [81], structural alignment with crystallized homologs revealed a strongly conserved T3SS fold (RMSD = 1.34 - 4.33 Å) despite low sequence identity (18-46%; Table 1), indicating that Ct has retained the ancestral injectisome architecture.

### 3.2 Construction of oligomeric assemblies

Oligomeric assemblies were built using three complementary approaches: AlphaFold 3, template-based modeling (SWISS-MODEL), and a Biopython-based script (Table 1). AlphaFold 3 reliably predicted only the CdsRST complex (ipTM 0.73) and failed on large or template-poor assemblies such as the CdsL dimer (ipTM 0.22), a known limitation for high-protomer complexes [62]. SWISS-MODEL was therefore used for CdsF, CdsC, and CdsV oligomers (Table 1), assessed by GMQE and QMEANDisCo Global scores (reliable at > 0.5 and > 0.4, respectively) [56,82]. CdsF scored GMQE and QMEANDisCo Global scores of 0.54 and 0.65 ± 0.05, respectively, indicating a reliable model, whereas CdsC (0.27 and 0.46 ± 0.05) and CdsV (0.26 and 0.52 ± 0.05) showed sub-threshold GMQE values, likely reflecting their low (< 30%) target-template identity (Table 1) [83]. Overall, the SWISS-MODEL assemblies remained reliable by QMEANDisCO Global scores. As SWISS-MODEL could not model the CdsD, CdsJ and CdsN oligomers, these were assembled with the Biopython-based script using homolog crystal structures as templates (Table 1).

### 3.3 Quality assessment of assembled subcomplexes

The subcomplex models were validated against crystal structures of their T3SS homologs (positive controls). In Ramachandran plot analysis (Ramplot), all subcomplexes showed 97.33-100% of residues in favored regions (Table S2, Figure S2), with the autoprotease, ATPase and stator proteins showing 100% of residues in the favoured region, confirming stereochemically reliable models. PRODIGY was used to predict the binding energetics between adjacent monomers. The needle complex, basal body (except CdsJ), and export apparatus exhibited binding energetics comparable to their homologs (Table 2). CdsJ and CdsN bound markedly more tightly than their homologs. CdsJ-CdsJ interaction in the IMR showed ΔG = −29.1 kcal/mol and K_d_ = 4.8×10^-22^ M (vs SctJ in *S. enterica*, ΔG = −8.5 kcal/mol, K_d_ = 5.5×10^-07^ M), and CdsN-CdsN interaction in the ATPase complex showed ΔG = −65.6 kcal/mol and K_d_ = 7.5×10^-49^ M (vs SctN in *E. coli*, ΔG = −12.4 kcal/mol, K_d_ = 8.3×10^-10^ M), indicating strong, thermodynamically stable interactions. PDBePISA further resolved the hydrogen bonds, salt bridges, and hydrophobic contacts at each interface, and the hydrogen bonding residues were mapped within each subcomplex (Figures 2-4; Table 2).

**Figure 2:**
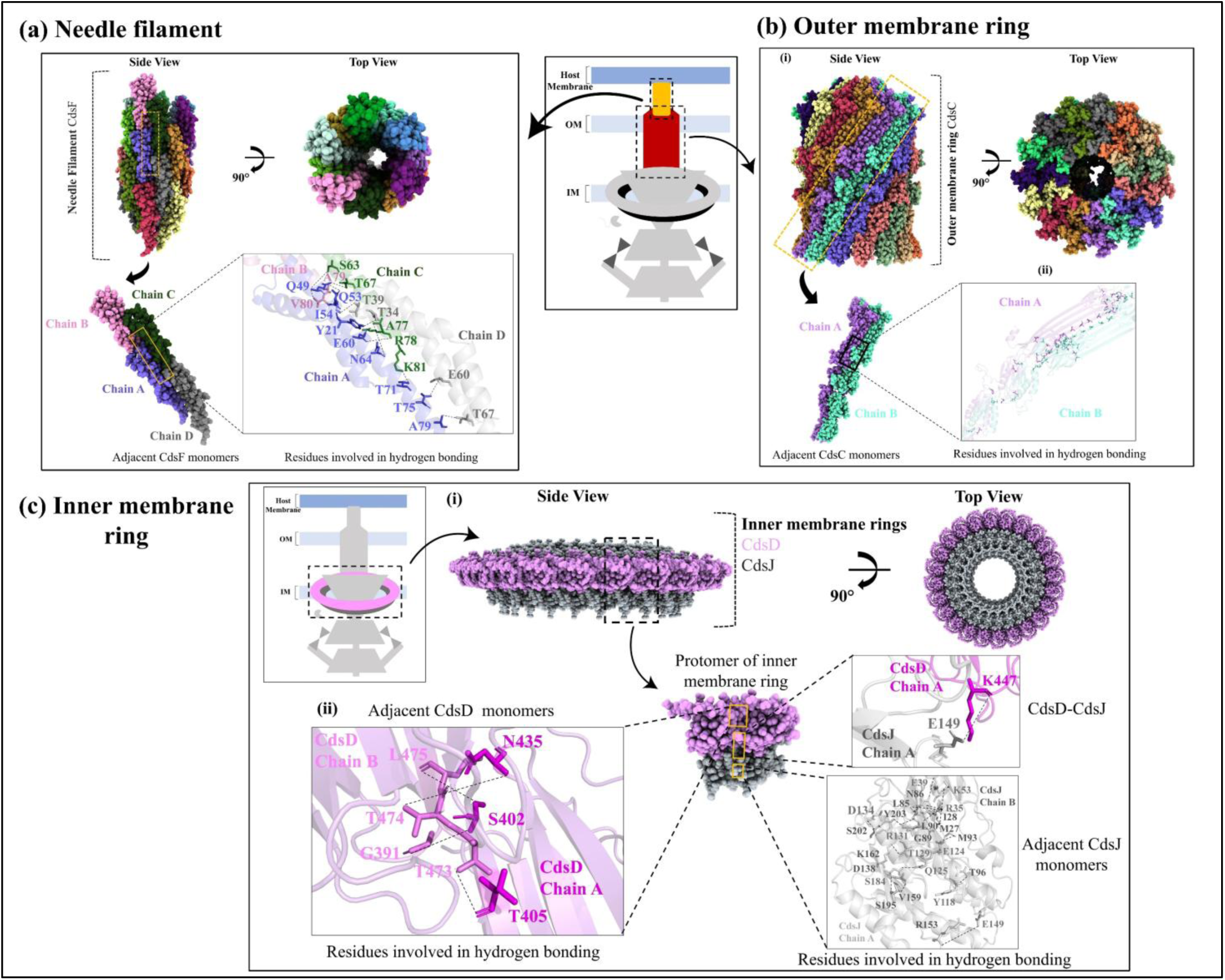
Structural arrangement of the needle filament, the outer membrane ring and inner membrane ring of Ct T3SS. Panel **(a)** illustrates the needle complex of T3SS. **(i)** Side and top views of the modeled needle complex are shown highlighting its spiral-like arrangement. **(ii)** The key amino acid residues involved in hydrogen bond between adjacent CdsF monomers are shown. Panel **(b)** depicts the outer membrane ring of the T3SS. **(i)** Side and top views of the outer membrane ring formed by CdsC are shown, which demonstrates its hollow and cylindrical architecture. **(ii)** The amino acid residues participating in hydrogen bond between adjacent CdsC monomers are illustrated. Panel **(c)** represents the inner membrane ring formed by CdsD and CdsJ. An enlarged view of a single protomer comprising CdsD (magenta) and CdsJ (grey) is shown. **(i)** The side view displays a solid ring-like arrangement of the complex, while the top view reveals a 48-fold symmetrical assembly surrounding a central channel. **(ii)** The key amino acid residues involved in hydrogen bond between adjacent CdsD, CdsJ, and CdsD-CdsJ monomers are highlighted.

**Table 2:**
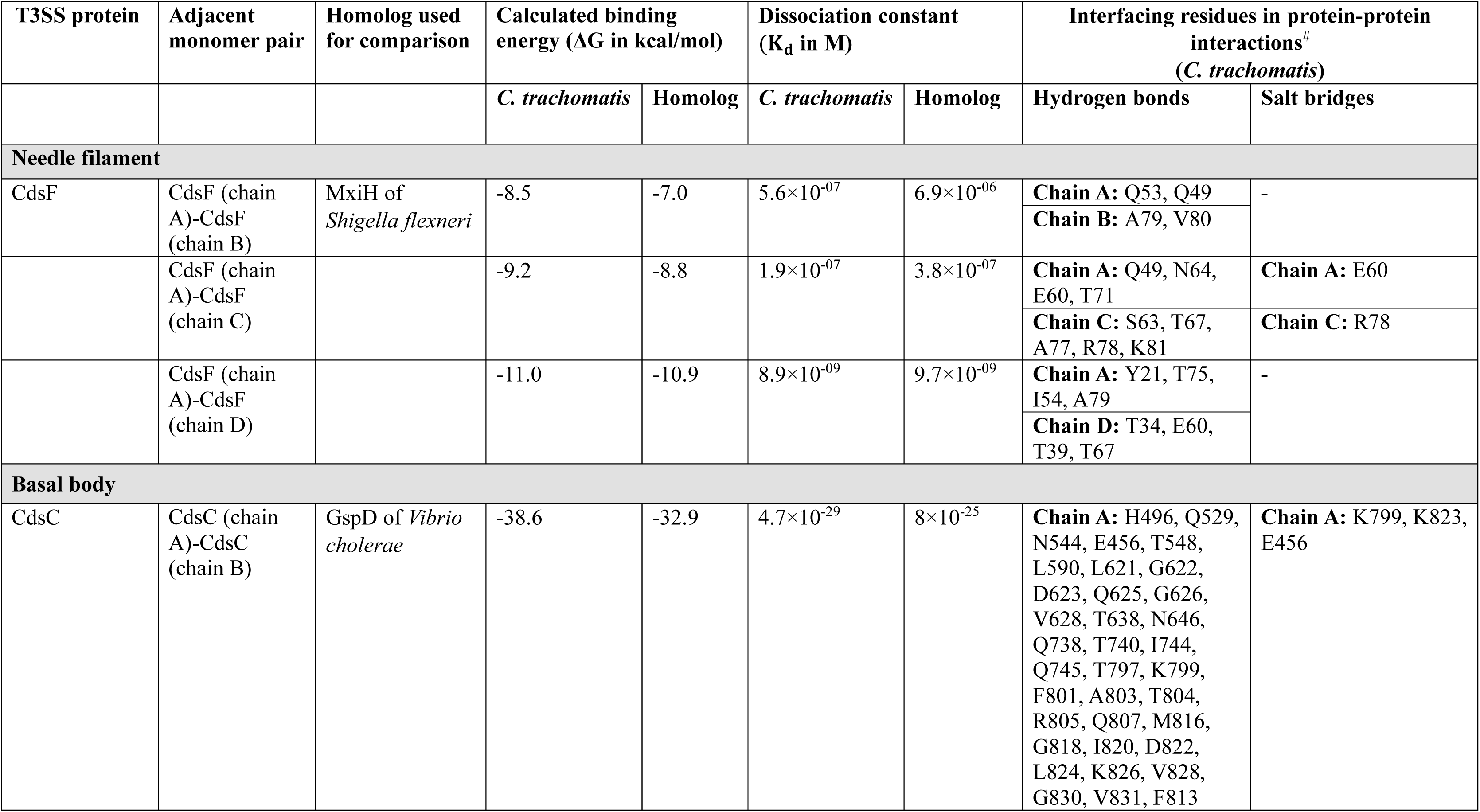
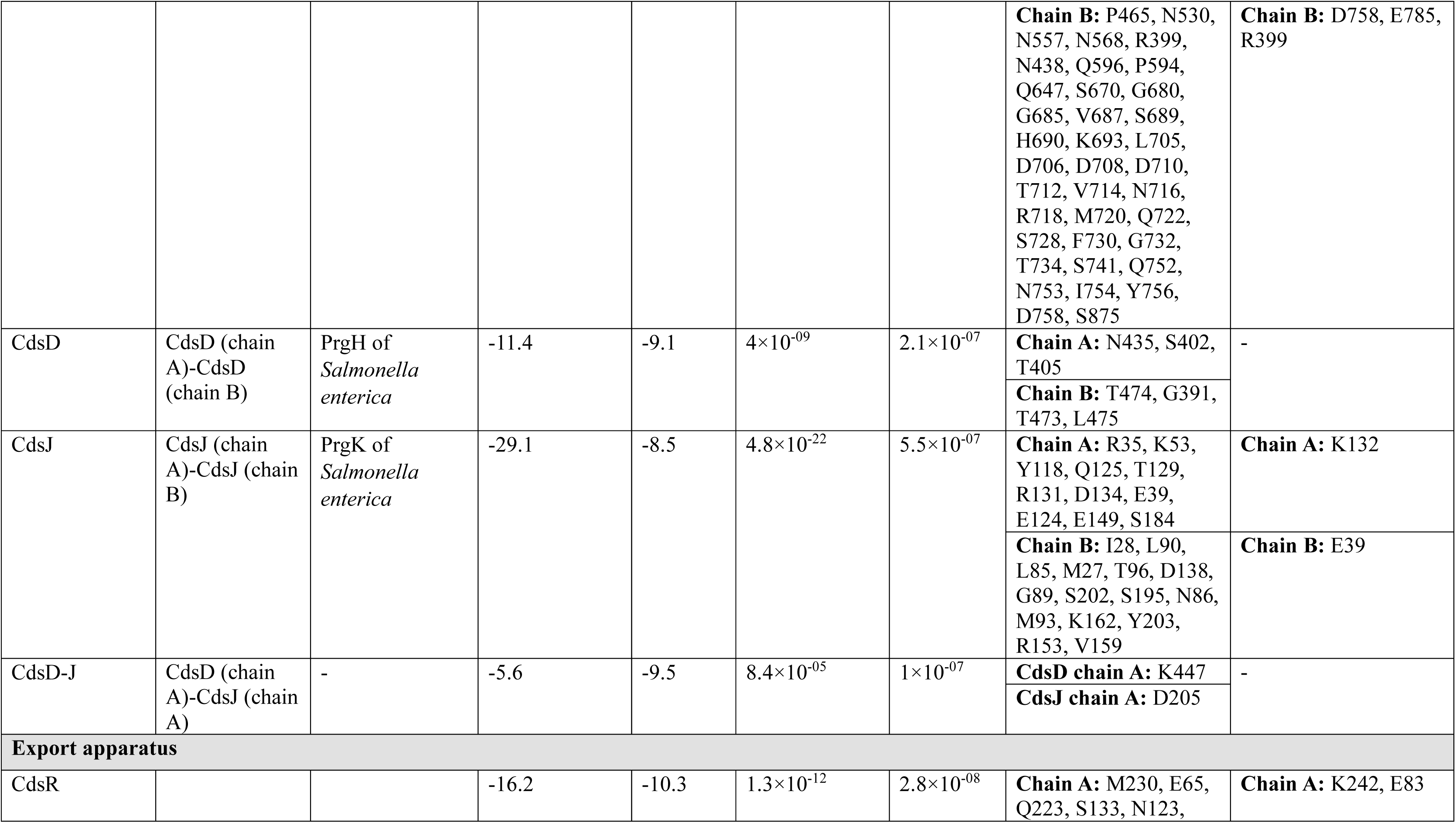
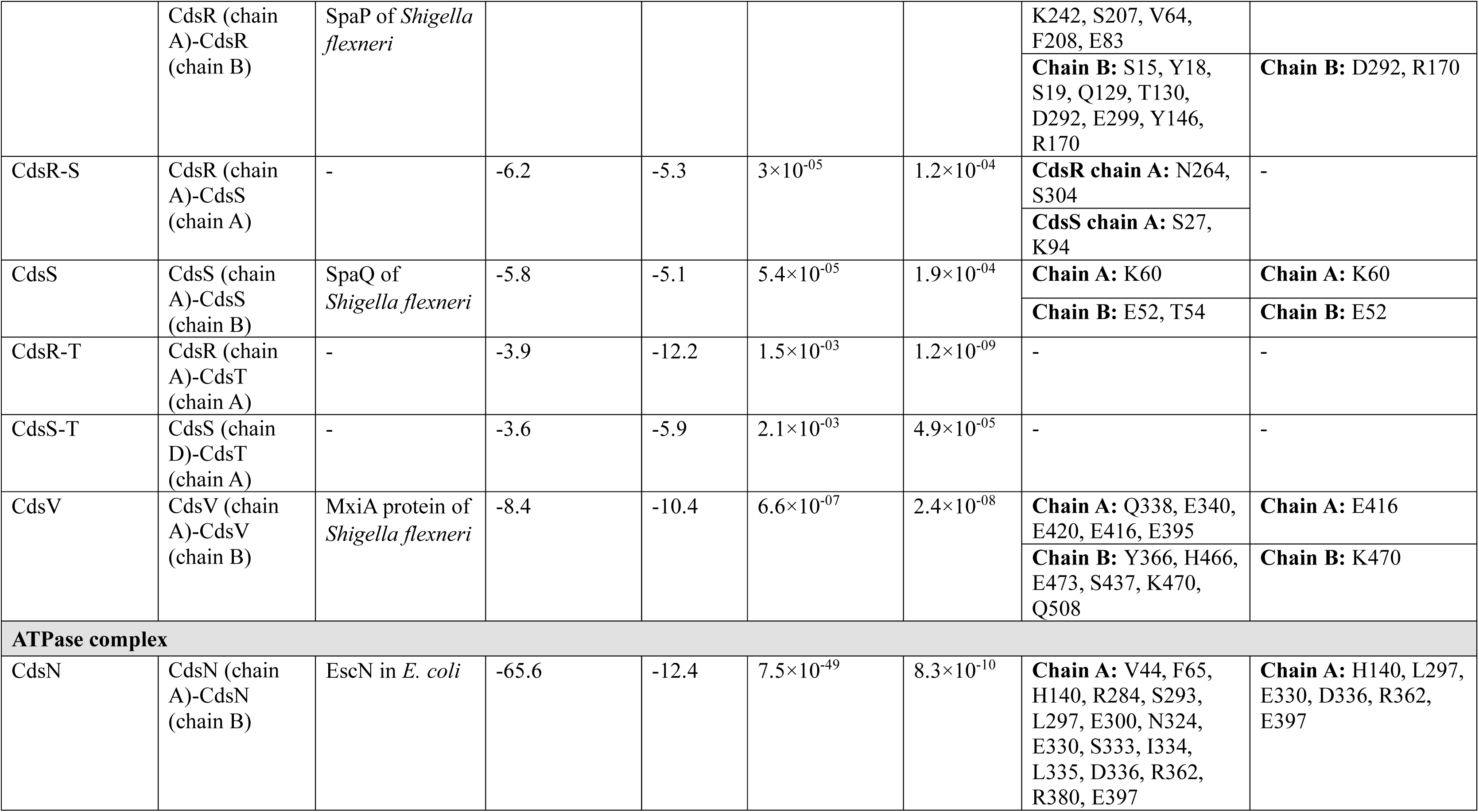
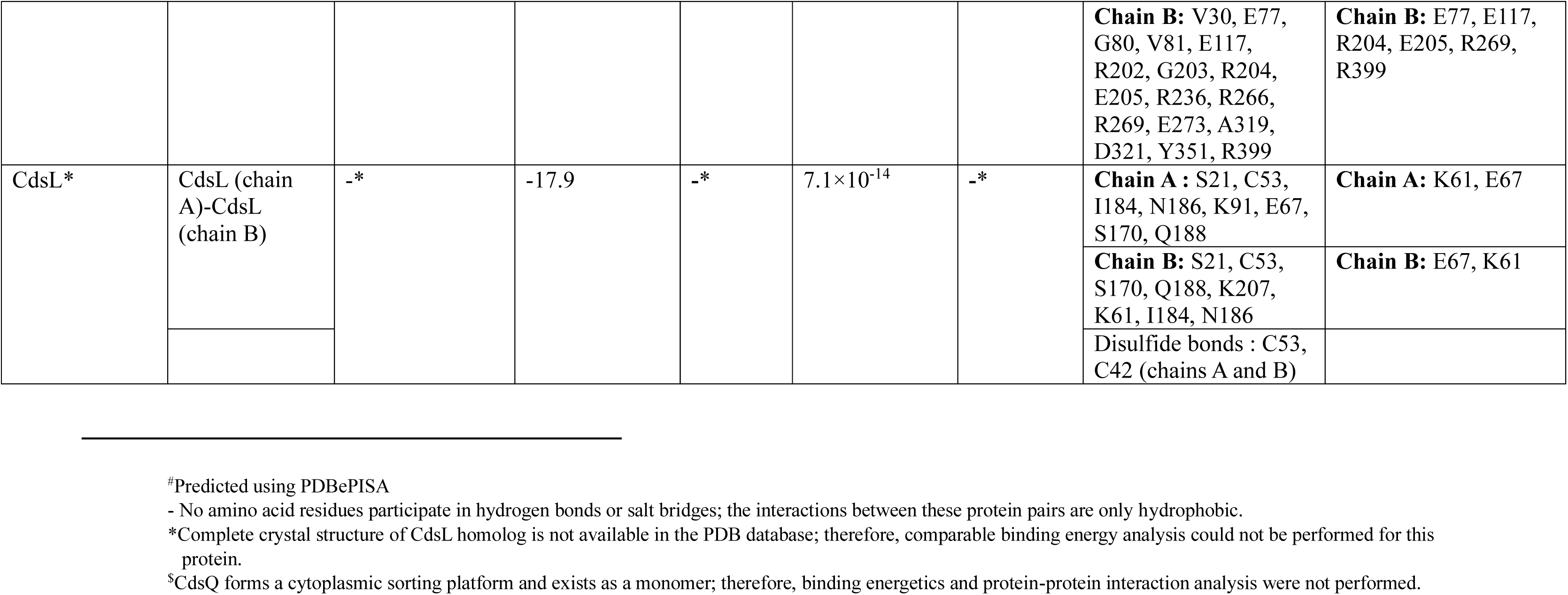
Binding affinity and protein-protein interaction analysis of T3SS subcomplexes^$^ in *C. trachomatis* and their structural homologs.

### 3.4 Assembly of the complete T3SS complex

After validation, the subcomplexes were manually assembled into the complete Ct T3SS (Figure 5), guided by the predicted model and the T3SS organization of *S. enterica* and *S. flexneri* [14,26]. The needle complex, composed of 23 copies of CdsF in a right-handed helix, forms the hollow conduit for effector injection (Figure 2a). Beneath it, CdsC forms a symmetric pentadecameric OMR that anchors the apparatus in the OM and gates substrate passage into the periplasm. Its C-terminal domains form an OM barrel, and its N-terminal domains extend into the periplasm to contact the IMR (Figure 2b). Within the IMR, CdsD and CdsJ each adopt 24-fold symmetry and form two nested concentric rings, CdsD enveloping CdsJ (Figure 2c). PPM analysis (with homolog as the positive control) showed OMR membrane orientation to be highly consistent between species (Figure S3a and S3b). PPM could not model IMR insertion for either *S. enterica* or Ct (Figure S3c and S3d), likely a server limitation. The IMR surrounds the export apparatus-the CdsRST complex, export gate CdsV, and autoprotease CdsU. The CdsRST complex model showed a 5:4:1 hetero-oligomeric stoichiometry as in *S. flexneri* [32,33]: five CdsR subunits form the outer ring-like scaffold, four CdsS subunits sit asymmetrically beneath, and a single CdsT occupies the central cavity, giving a conical, membrane-embedded architecture (Figure 3a and 3b). CdsV forms a homononameric export gate below CdsRST (Figure 3c, 3d and 5), and monomeric autoprotease CdsU lies near the CdsRST complex (Figure 3e and 5). In the ATPase complex, CdsN assembles into a homohexameric ring that powers substrate unfolding and extraction (Figure 4a and 4b), positioned below the export apparatus and linked to the export gate via the stalk protein CdsO. As CdsO is already crystallized (PDB ID: 3K29), it was used only in the final assembly. CdsL functions as a stator whose dimer associates with a CdsN monomer (Figure 4c and 4d) [45,84], connecting CdsN to the C-ring protein CdsQ, which forms a pod-like structure at the end of each CdsL dimer (Figure 4e).

**Figure 3:**
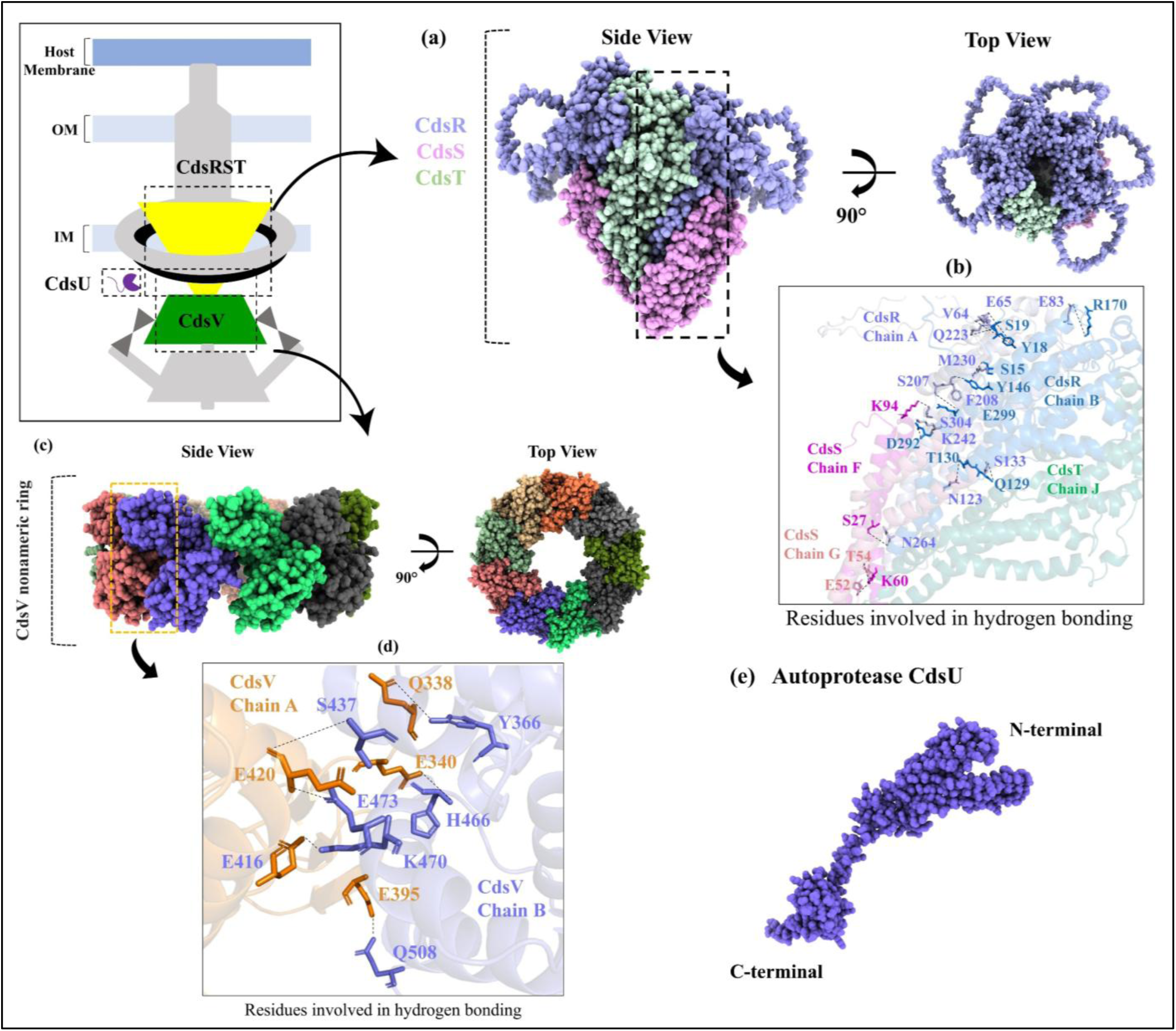
Structural arrangement of the Ct T3SS export apparatus. **(a)** Side and top views of the hetero-oligomeric CdsRST complex are shown, which demonstrates its conical architecture. **(b)** Enlarged view showing key amino acid residues involved in hydrogen bond between adjacent CdsR, CdsS, and CdsT monomers. **(c)** Side and top views of the homononameric CdsV export gate. **(d)** The amino-acid residues involved in hydrogen bond between adjacent CdsV monomers are shown. **(e)** Structural model of the autoprotease CdsU, with the N-terminal and C-terminal regions indicated.

**Figure 4:**
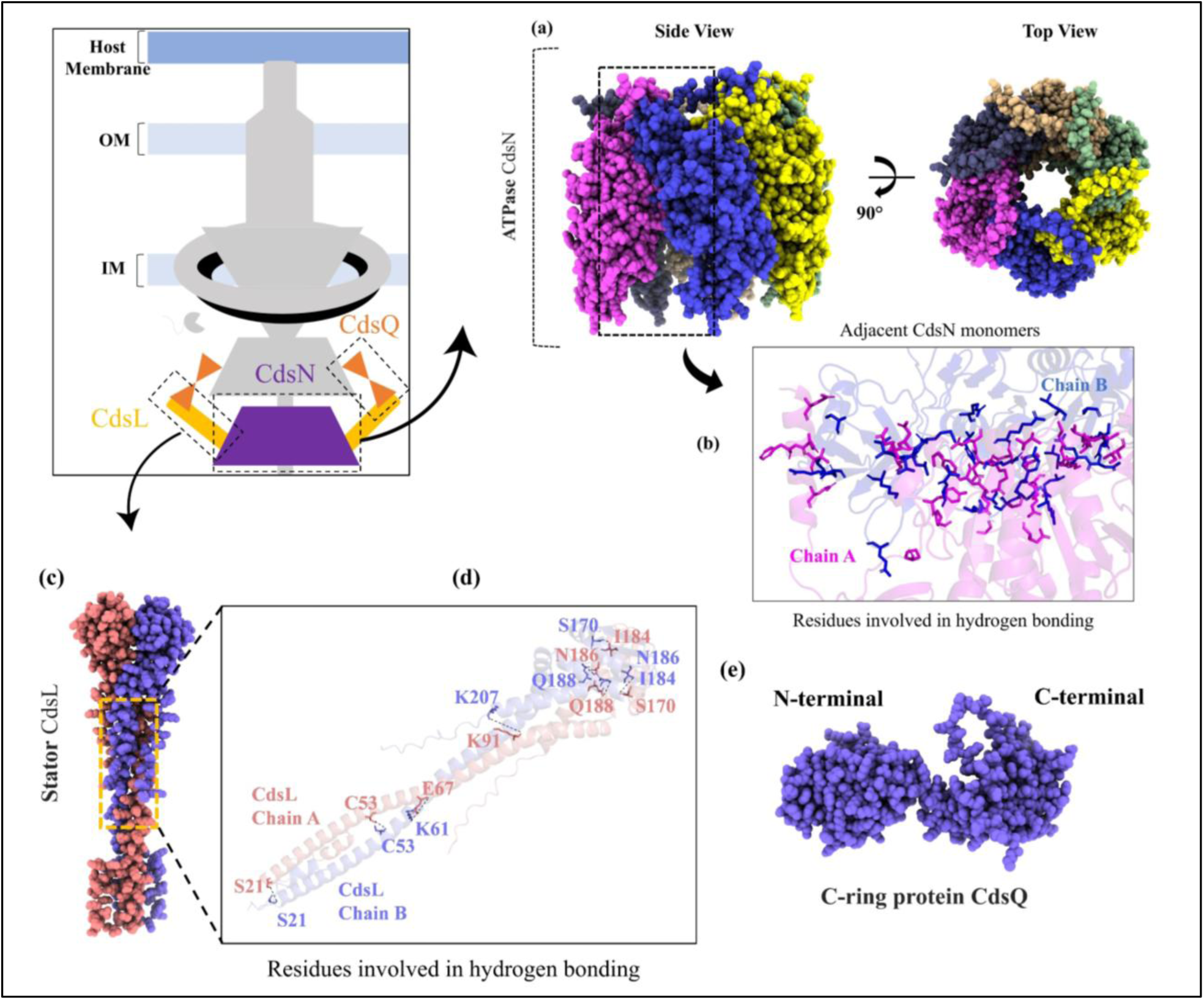
Structural arrangement of the Ct T3SS ATPase complex and C-ring component. **(a)** Side and top views of the hexameric CdsN ATPase are shown. **(b)** Enlarged view highlighting key amino acid residues involved in hydrogen bond between adjacent CdsN monomers. **(c)** Structural model of the CdsL dimer stator protein is shown. **(d)** The key amino acid residues involved in hydrogen bond between adjacent CdsL monomers are shown. **(e)** Structural model of the C-ring component CdsQ, with the N-terminal and C-terminal regions indicated.

**Figure 5:**
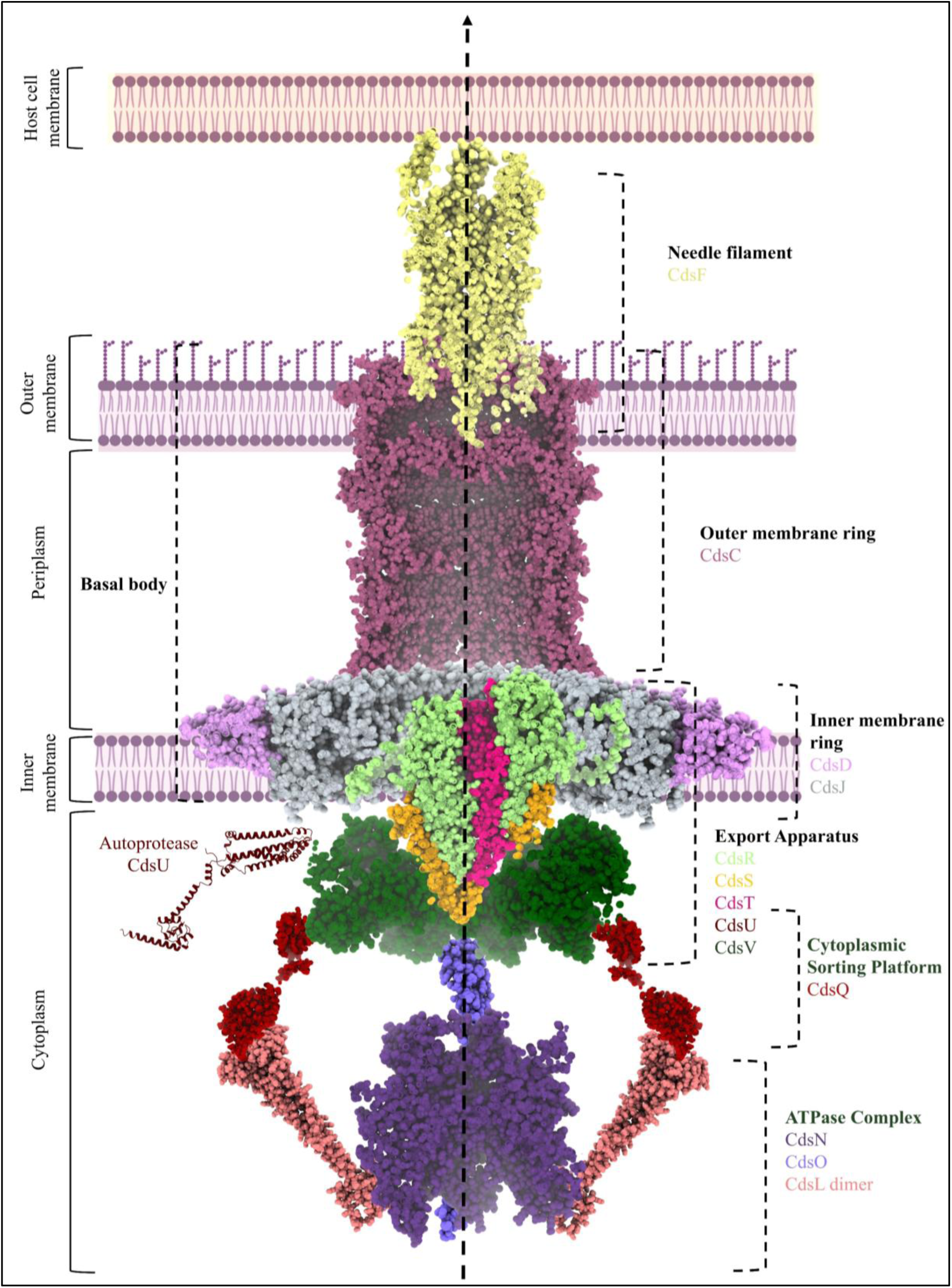
Complete architecture of the Ct T3SS complex. The overall structure of the Ct T3SS complex can be divided into five subcomplexes: needle complex (CdsF); basal body (CdsC, CdsD, and CdsJ); export apparatus (CdsR, CdsS, CdsT, CdsU, and CdsV); ATPase complex (CdsN, CdsL, and CdsO); and cytoplasmic sorting platform (CdsQ). The basal body of the apparatus spans the inner membrane (IM), periplasmic space, and OM of the bacteria.

### 3.5 Homology modeling and model quality assessment of CdsN ATPase

Although CdsN assembles into a homohexameric ring within the intact T3SS (Figure 6a), the present drug-repurposing analysis targets the PPI interface between adjacent subunits; we therefore modeled two adjacent CdsN monomers, and the term ‘dimer’ is used throughout the drug repurposing sections to denote this pair of adjacent monomers representing the oligomerization interface. The CdsN dimer modeled with PPI3D is shown in Figure 6. Chain A contains four N-terminal β-strands and chain B has six β-strands, but their C-terminal domains are similar, each with three α-helices (Figure 6b and 6c). The two chains aligned at RMSD 0.91 Å, consistent with the *E. coli* EscN chains (RMSD 0.79 Å) and attributable to identical subunits adopting distinct conformations and catalytic states, as reported for EscN [85]. Superimposition onto the EscN dimer (PDB ID: 6NJP) gave an RMSD of 1.17 Å, supporting model reliability (Figure S4). ProSA-web yielded a Z-score of −9.75, consistent with experimental structures (Figure S5a), and Ramachandran plot placed 95.355% of residues in favored regions, 3.178% in allowed regions, and 1.467% in disallowed regions (Figure S5b), confirming a good quality model. PRODIGY predicted ΔG = −16.5 kcal/mol and K_d_ = 8.2 × 10^-^¹³ M for the CdsN dimer. Both ΔG and K_d_ were more favorable than the *E. coli* EscN control (ΔG = −13.7 kcal/mol, K_d_ = 8.8×10^-11^ M), indicating strong and stable self-association. Sequence comparison with the human proteome revealed no significant homology (identities < 35%), supporting CdsN as a drug target.

**Figure 6:**
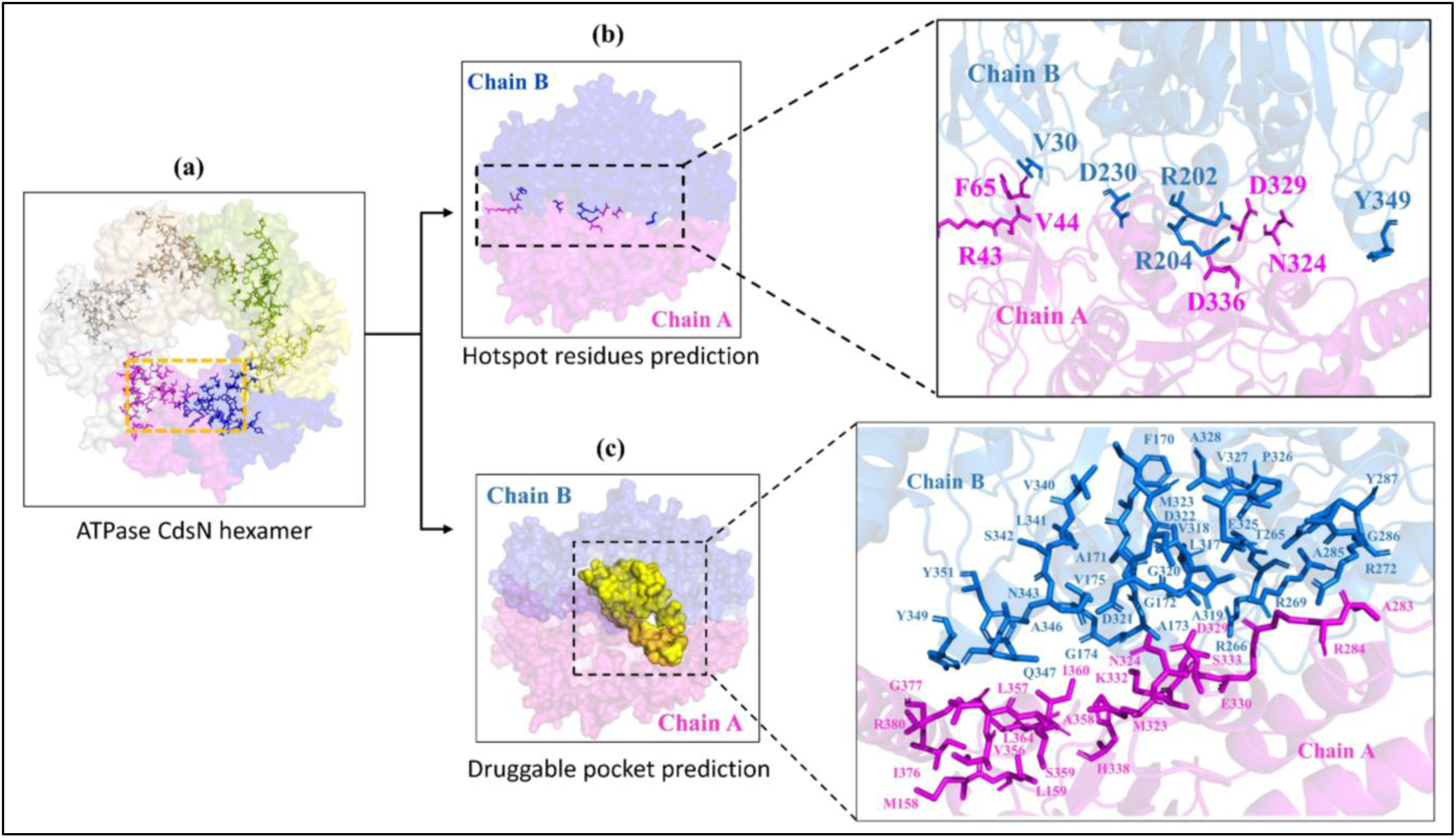
Prediction of hotspot residues and druggable pocket. **(a)** Schematic representation of the CdsN hexamer oligomerization interface used for structure-based virtual screening. **(b)** DrugScore PPI analysis predicted six interface hotspot residues in chain A and five in chain B, as shown in the right panel. **(c)** The druggable pocket predicted by DOGSiteScorer is shown, with the corresponding amino acid residues forming the pocket displayed in the right-side panel.

### 3.6 Prediction of hotspot residues and druggable pocket

Hotspot residues contribute disproportionately to interface binding free energy and are essential for complex stability [86]. DrugScore PPI predicted six hotspot residues in chain A (R43, V44, F65, N324, D329 and D336) and five in chain B (V30, R202, R204, D230 and Y349) (Figure 6b), marking them as promising targets for small-molecule inhibition. DOGSiteScorer identified 30 potential druggable pockets at the CdsN oligomerization interface, of which the highest-scoring (druggability score 0.87, close to 1.0) was selected (Figure 6c). Structure-based virtual screening was then performed against both the hotspot region and this druggable pocket.

### 3.7 Virtual screening of compounds

Two databases were screened: e-Drug3D (approved drugs and commercially available drug fragments) [75], and IMPPAT 2.0 (phytochemicals from Indian medicinal plants, encompassing data on 4,010 Indian medicinal plants, 17,967 phytochemicals and 1,095 therapeutic uses) [76]. Screening against the hotspot residues yielded 40 e-Drug3D and two IMPPAT hits with docking scores ≤ −7.0 kcal/mol (Tables S3 and S5), a threshold generally indicative of good affinity between protein and ligand [77]. Screening against the druggable pocket gave 23 e-Drug3D and seven IMPPAT hits (Tables S4 and S6). All hits with docking score ≤ −7.0 kcal/mol were advanced to ADMET screening.

### 3.8 ADMET screening

The 72 hits from both databases targeting the two binding regions were subjected to ADMET evaluation using QikProp against eight pharmacokinetic criteria: one violation of Lipinski’s Rule of Five, ≥80% predicted human oral absorption, QPPCaco > 500, QPPMDCK (<25 poor, 25-500 moderate and >500 excellent), QPlogHERG < −5, donor HB ≤ 5, acceptor HB ≤ 10, and QPlogPo/w (−2 to 6.5) [78] (Tables S3-S6). Three e-Drug3D compounds-M Roflumilast (Drug ID: 1537), Elacestrant (Drug ID: 2081), and Tecovirimat (Drug ID: 1889)-met the criteria (Table 3), whereas no IMPPAT phytochemicals qualified, most failing on predicted membrane permeability (QPPMDCK and QPPCaco) and intestinal absorption (Tables S5, S6).

**Table 3:** Selected parameters, including docking scores, binding free energies, and pharmacokinetic properties of selected drug candidates targeting the CdsN oligomerization interface.

| Selected Parameters | Drug candidates |  |  |
| --- | --- | --- | --- |
|  | M Roflumilast | Elacestrant | Tecovirimat |
| Docking Score ($> -7.0$ kcal/mol) | -7.165 | -7.118 | -7.127 |
| $\Delta G_{\text{bind}}^{\#}$ | -39.76 | -67.98 | -51.42 |
| SASA<sup>a</sup> | 648.514 | 776.187 | 605.725 |
| QPPMDCK<sup>b</sup> ($< 25$ poor; 25-500 moderate; $> 500$ excellent) | 8784.547 | 361.172 | 1812.978 |
| QPPCaco<sup>c</sup> ($> 500$) | 1605.376 | 680.758 | 849.205 |
| QPlogPo/w<sup>d</sup> ($-2$ to $6.5$) | 5.004 | 6.391 | 3.9 |
| donorHB ($\leq 5$) | 1 | 2 | 0.25 |
| acceptHB ($\leq 10$) | 4.25 | 4 | 4.75 |
| QPlogHERG<sup>e</sup> ($< -5$ = safer) | -5.424 | -6.229 | -5.167 |
| Rule of five (Lipinski rule) ($\leq 1$) | 1 | 1 | 0 |
| Percentage of Human Oral Absorption ($> 80\%$ = good) | 100 | 100 | 100 |

<sup>#</sup>Estimation of $\Delta G_{\text{bind}}$ (free energy of binding, kcal/mol) using MM-GBSA analysis.
<sup>a</sup>SASA (solvent-accessible surface area) describes the extent of a molecular surface exposed to the solvent.
<sup>b</sup>QPPMDCK (nm/s) predicts cellular membrane permeability and estimates the ability of drug molecules to penetrate cell membrane.
<sup>c</sup>QPPCaco (nm/s) defines the predicted Caco-2 cell permeability, which indicates intestinal absorption potential.
<sup>d</sup>QPlogPo/w represents the predicted water partition coefficient and reflects the compound lipophilicity.
<sup>e</sup>QPlogHERG is the predicted potential for hERG channel inhibition, which is associated with cardiac toxicity risk.

M Roflumilast (a derivative of the FDA-approved PDE4 inhibitor [87]) and Elacestrant (a selective estrogen receptor degrader [88]) were identified against the hotspot residues. Both satisfied Lipinski’s rule (one permissible violation), with docking scores of −7.165 and −7. 118, respectively (Table 3; Figure 7a, 7b) and showed 100% human oral absorption and strong intestinal absorption with QPPCaco of 1605.376 and 680.758 nm/s, respectively. M Roflumilast showed high predicted membrane permeability (QPPMDCK = 8784.547 nm/s)-an important property for targeting intracellular pathogen. Elacestrant showed moderate membrane permeability (QPPMDCK = 361.172 nm/s), warranting experimental validation. Their lipophilicity (QPlogPo/w = 5.004 for M roflumilast, 6.391 for Elacestrant; ideal −2.0 to 6.5) indicated favorable physicochemical properties for membrane permeation. Both compounds also showed acceptable QPlogHERG of −5.424 and −6.229, respectively (ideal < −5), indicating lower risk of hERG channel inhibition. Tecovirimat, an FDA-approved antiviral [89], was identified against the druggable pocket (Table 3, Figure 7c). It fully complied with Lipinski’s rule of five (no violations), with 100% human oral absorption, QPPCaco of 849.205 nm/s, QPPMDCK of 1812.978 nm/s, QPlogPo/w of 3.9, and QPlogHERG of −5.167. These three compounds were considered further for binding energetics evaluation and MD simulation.

**Figure 7:**
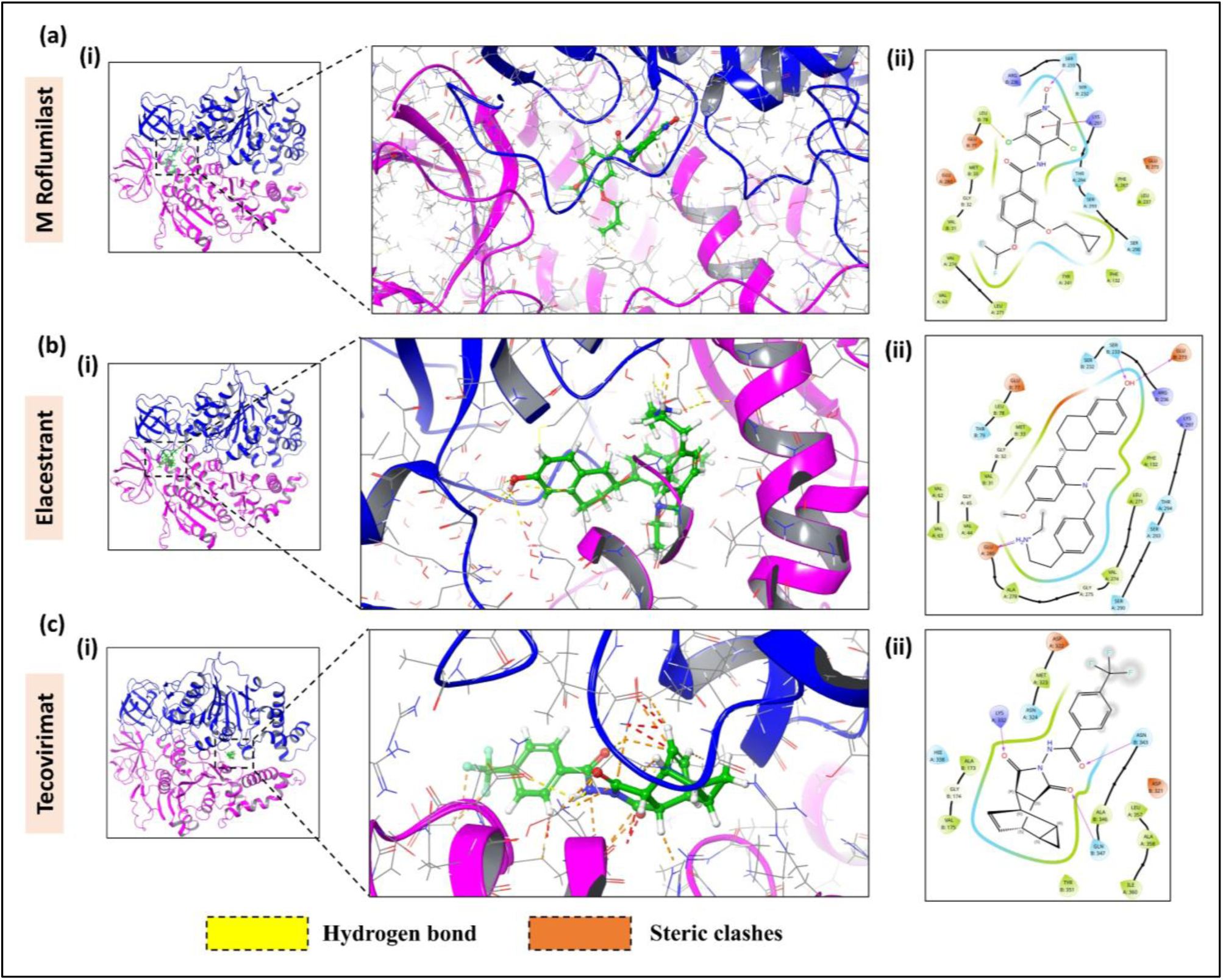
Molecular docking of the CdsN dimer with the eDrug3D compound library. Panels **(a-c)** correspond to the virtual screening of the eDrug3D compounds (M Roflumilast, Elacestrant and Tecovirimat) against the predicted hotspot residues and druggable pocket at the CdsN dimeric interface. For each complex: **(i)** The CdsN dimer-ligand complex is shown docked. **(ii)** The binding interactions between the CdsN dimer and ligand are shown, with magenta arrows denoting hydrogen bond interactions between the protein and ligand.

### 3.9 Binding free energies of the docked complexes

Prime/MM-GBSA showed that all three ligands formed energetically favorable complexes with the CdsN dimer, with Elacestrant exhibiting strongest binding affinity (ΔG_bind_ = −67.98 kcal/mol), driven by van der Waals (ΔG_vdW_ = −39.8 kcal/mol), hydrophobic interactions (ΔG_lipo_ = −32.98 kcal/mol), and a favorable electrostatic contribution (ΔG_cou_ = −16. 34 kcal/mol) (Tables 3, S7).

M Roflumilast bound stably (ΔG_bind_ = −39.76 kcal/mol), mainly via van der Waals (ΔG_vdW_ = −41.3 kcal/mol) and hydrophobic (ΔG_lipo_ = −15.42 kcal/mol) contributions, with minimal Coulombic (ΔG_cou_ = 6.46 kcal/mol) and covalent energy (ΔG_cov_ = 1.06 kcal/mol) (Tables 3, S7). Tecovirimat gave ΔG_bind_ = −51.42 kcal/mol, with substantial van der Waals (ΔG_vdW_ = −39.8 kcal/mol) and electrostatic (ΔG_coul_ = −42.43 kcal/mol) contributions and further hydrophobic stabilization (ΔG_lipo_ = −12.88 kcal/mol) (Tables 3, S7). Thus, although docking scores were close to the −7.0 kcal/mol threshold, MM/GBSA analysis confirmed thermodynamically stable complexes for all three ligands, with Elacestrant as the most favorable.

### 3.10 MD simulation

For each protein-ligand complex, conformational stability was primarily assessed using four iterative 50 ns simulations, since multiple short trajectories improve conformational sampling relative to a single long run [90]. This resulted in a cumulative simulation time of 200 ns per system [49]. As a backup, a single continuous 100 ns simulation was also performed for each complex to independently confirm these trends.

#### CdsN dimer-M Roflumilast complex

Across the four iterative 50 ns simulations, protein backbone and ligand RMSDs (∼2.73 and ∼3.25 Å, respectively) initially increased during the first continuation simulation run before decreasing to ∼1.68 and ∼1.73 Å by the end of the fourth simulation (Table 4 and Figure 8a), suggesting an initial conformational rearrangement of the ligand within the binding pocket followed by convergence to a stable binding pose. This trend was consistent with the 100 ns run, which showed average protein backbone and ligand RMSDs of 2.8 and 3.0 Å, respectively, with the ligand fluctuating more than the protein but without evident dissociation (Figures 8a and S6a). RMSF values in the 100 ns run were low (0.5 to 1.5 Å) for ligand-interacting residues involved in hydrogen bonds (chain A: Glu-280; and chain B: Ser-233, Met-33) and the ligand remained compact (rGyr; ∼ 4.75-4.85 Å; SASA ∼80 −90 Å^2^) (Figure S6a, S7a).

**Figure 8:**
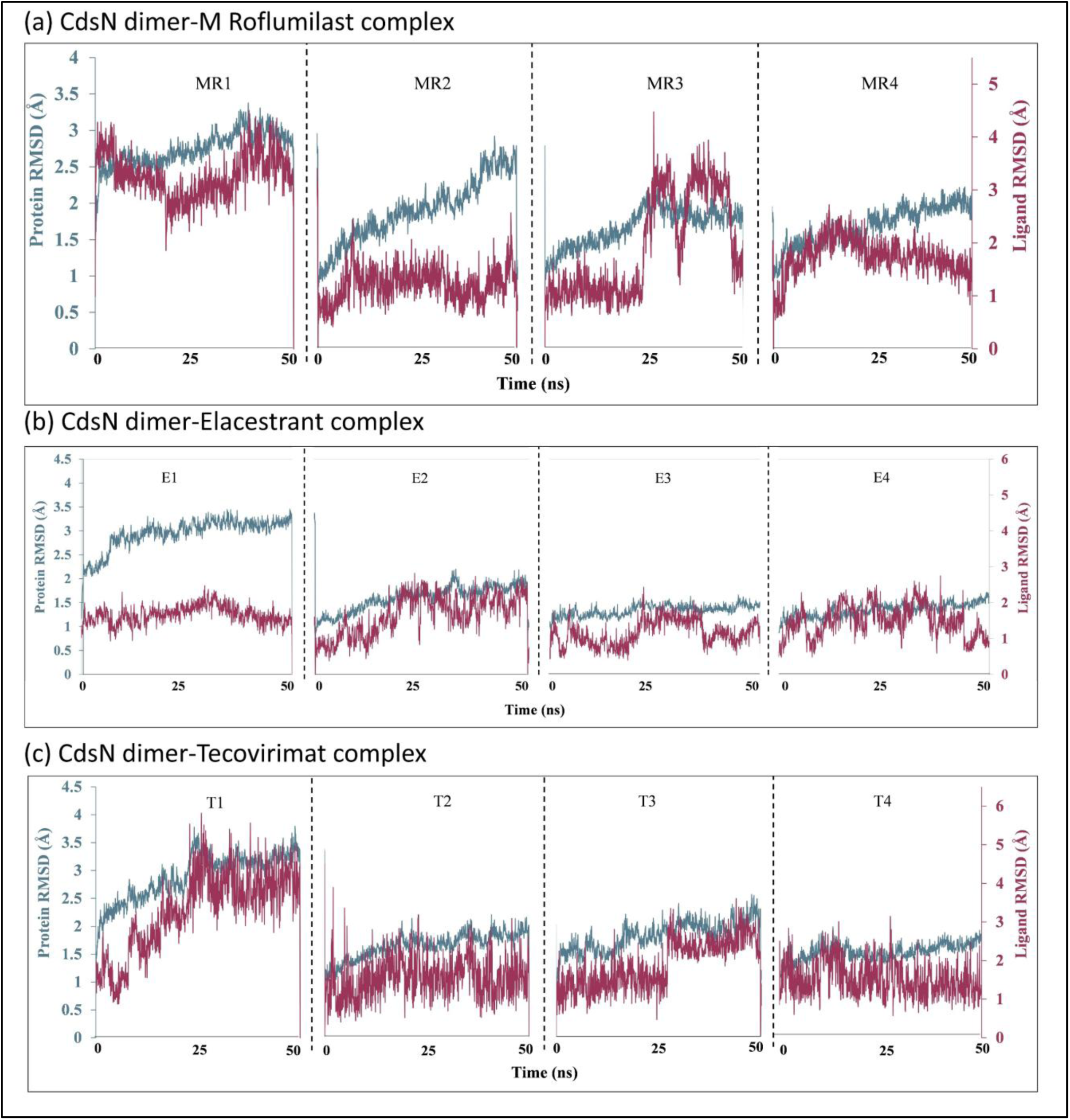
Comparative RMSD profiles of protein-ligand complexes. MD simulation of CdsN dimer with **(a)** M roflumilast, **(b)** Elacestrant, and **(c)** Tecovirimat, over iterative 50 ns time scale. MR1-MR4, E1-E4, and T1-T4 represent the first to fourth 50 ns continuation simulations of the CdsN dimer-M Roflumilast, CdsN dimer-Elacestrant, and CdsN dimer-Tecovirimat complexes, respectively.

**Table 4:** Average RMSD values of protein (CdsN dimer) and ligand during 50 ns MD Simulation.

| Protein-Ligand Complex | Simulation Run<sup>#</sup> | Average Protein Backbone RMSD (Å) | Average Ligand RMSD with respect to Protein (Å) |
| --- | --- | --- | --- |
| CdsN dimer - M Roflumilast | MR1 | 2.73 | 3.25 |
|  | MR2 | 1.89 | 1.21 |
|  | MR3 | 1.67 | 1.93 |
|  | MR4 | 1.68 | 1.73 |
| CdsN dimer - Elacestrant | E1 | 2.93 | 1.66 |
|  | E2 | 1.60 | 1.65 |
|  | E3 | 1.33 | 1.15 |
|  | E4 | 1.34 | 1.47 |
| CdsN dimer - Tecovirimat | T1 | 2.90 | 3.28 |
|  | T2 | 1.65 | 1.53 |
|  | T3 | 1.82 | 1.88 |
|  | T4 | 1.53 | 1.51 |

<sup>#</sup>MR1-MR4, E1-E4, and T1-T4 represent the first to fourth 50 ns continuation simulations of the CdsN dimer-M Roflumilast, CdsN dimer-Elacestrant, and CdsN dimer-Tecovirimat complexes, respectively.

#### CdsN dimer-Elacestrant complex

Across the four 50 ns continuation simulations, both protein backbone and ligand RMSDs decreased progressively (2.9 to 1.3 Å and 1.6 to 1.4 Å, respectively), indicating that Elacestrant achieved a stable accommodation within the CdsN pocket with minimal positional rearrangement (Table 4 and Figure 8b). This stability was corroborated by the 100 ns run, in which the average protein backbone and ligand RMSDs (∼3.0 and 1.9 Å, respectively) reflected a structurally stable protein undergoing modest ligand-induced adjustment (Figure S6b). RMSF values for ligand-interacting residues (hydrogen bonds-chain A: Val-63, Ser-293; and chain B: Val-31) were low (0.6-1.2 Å) (Figure S6b), and the ligand showed stable compactness (rGyr: 4.35-4.65 Å) with a relatively consistent degree of solvent exposure (SASA ∼10-45 Å^2^) (Figure S7b).

#### CdsN dimer-Tecovirimat complex

During the first 50 ns simulation, both protein and ligand RMSDs increased (2.90 and 3.28 Å, respectively), indicating an initial conformational rearrangement within the binding pocket, and a progressive decrease over the subsequent three simulation runs indicated that Tecovirimat gradually adopted a stable binding orientation without dissociation (Table 4 and Figure 8c). This was consistent with the 100 ns run, which showed average protein backbone and ligand RMSDs of 3.1 and 3.8 Å, respectively, with no evidence of ligand dissociation (Figure S6c). RMSF values for ligand-interacting residues (hydrogen bonds-chain A: Asp-322, Lys-332; and chain B: Ala-173 and Gln-347) were low (1.0-2.0 Å), and the ligand remained compact (rGyr: ∼4.80 Å), though SASA (180-200 Å²) indicated that a substantial portion remained solvent-accessible (Figure S7c).

Overall, the iterative 50 ns simulations demonstrated that M Roflumilast, Elacestrant, and Tecovirimat all converged toward stable binding poses within the CdsN dimer, a conclusion corroborated by the 100 ns runs, through acceptable RMSD and RMSF profiles, persistent key intermolecular interactions, stable ligand compactness, and minimal variation in solvent exposure. Among the three, Elacestrant displayed the most favorable dynamic profile with the lowest RMSD values and the least structural fluctuation across both the iterative and backup simulations. This was consistent with its MM-GBSA results, which showed the most negative ΔG_bind_. Together, these dynamics support the potential of all three compounds as promising candidates for further experimental validation as repurposed therapeutics against Ct infection.

## 4. Discussion

Ct is an obligate intracellular pathogen in which the T3SS serves as a major virulence apparatus, delivering effector proteins such as TarP, TmeA, TmeB, and Incs directly from the bacterial cytosol into the host cell to establish infection and sustain intracellular replication and survival. Although the individual T3SS constituent proteins have been catalogued, the molecular-level architecture of the complete Ct T3SS has remained undefined—indeed, no complete T3SS structure has yet been fully resolved in any Gram-negative bacterium owing to the technical difficulty of characterizing such large multiprotein assemblies. Using an integrated computational pipeline [49], the present study delineates the first molecular-level model of the complete Ct T3SS and leverages it for structure-based antivirulence drug discovery.

We first identified and validated all 13 T3SS constituent proteins, combining literature curation with TXSSScan annotation and reverse-BLAST confirmation against the Ct D/UW-3/CX proteome. Notably, TXSSScan recovered only 8 of the 13 proteins; because it relies on conserved Hidden Markov Model profiles, the sequence divergence and dispersed genomic organization of the Ct T3SS—whose genes are distributed across four clusters forming a “pathogenicity archipelago” [21] rather than a single contiguous operon—likely limit its sensitivity, underscoring the value of the complementary reverse-BLAST step. The proteins were then modelled and assembled into five subcomplexes using three complementary approaches (AlphaFold 3, template-based modeling with SWISS-MODEL, and a Biopython-based script). For the three low pTM monomers (CdsF, CdsC, CdsD), cross-validation with four independent predictors supported the CdsF fold, whereas the higher RMSD values for CdsC and CdsD were traced to extensive intrinsically disordered regions that are inherently modeled differently across algorithms. Despite low overall sequence identity with homologs (18-46 %), structural alignment revealed a highly conserved architecture (low RMSD; Table 1), indicating that Ct has retained the ancestral injectisome fold even as its sequences diverged. The assemblies were further validated stereochemically (> 90% of residues in favored Ramachandran regions) and energetically, with PRODIGY-predicted ΔG and K_d_ comparable to those of crystallographic homolog controls.

Although the overall architecture and assembly energetics of the Ct T3SS resemble those of homologs from other Gram-negative bacteria, notable differences emerged. Two interfaces, CdsJ-CdsJ and CdsN-CdsN, bound markedly more tightly than their homologs (ΔG = −29.1 and −65.6 kcal/mol versus −8.5 and −12.4 kcal/mol). As these subunits form the IMR and the load-bearing ATPase hexamer, which must withstand the mechanical stress of substrate unfolding and injection, stronger self-association is mechanistically expected. We interpret this as an evolutionary adaptation favoring a stable, rapidly assembling injectisome rather than a modeling artifact, since the same pipeline reproduced the energetics of the homolog controls. These findings are consistent with other Ct-specific adaptations that reflect its obligate intracellular lifestyle. The T3SS is thought to have first evolved for basic survival and later been repurposed for pathogenesis [26]; the needle protein CdsF uniquely harbors cysteine residues that enable stage-dependent needle conformations [26]; in situ cryo-electron tomography indicates that host-membrane contact induces basal-body compaction [27]; and the periplasmic region of CdsD contains an extended α-helical segment, absent in other Gram-negative bacteria, proposed to support basal-body assembly and stability [41]. Together with developmentally regulated T3SS gene expression [92], these features distinguish the Ct injectisome despite its conserved core.

Because inhibition of T3SS function attenuates chlamydial replication and development [16,17], we exploited the assembled model for target selection, focusing on the oligomerization interface of the CdsN ATPase. The CdsN dimer model was of good quality (Z-score −9.75, and RMSD 1.17 Å to the *E. coli* EscN dimer) and self-associated strongly (ΔG = −16.5 kcal/mol), and DrugScorePPI and DOGSiteScorer defined a hotspot-rich, highly druggable pocket (druggability score 0.87) at this interface. Targeting oligomerization is mechanistically distinct from and complementary to earlier catalytic-site strategies in Ct and other pathogens [47,50,51]. Rather than competing with abundant intracellular ATP for the catalytic site, interface disruption is expected to prevent assembly of the functional hexameric ATPase, thereby stalling secretion [93]. The absence of significant CdsN homology with the human proteome (identity <35%) further supports its suitability as a selective antibacterial target. Structure-based virtual screening of the e-Drug3D and IMPPAT libraries against the interface, followed by ADMET filtering, yielded three FDA-approved candidates-M Roflumilast (Drug ID: 1537), Elacestrant (Drug ID: 2081), and Tecovirimat (Drug ID: 1889). Although their docking scores lay close to the −7.0 kcal/mol threshold, MM-GBSA analysis confirmed thermodynamically stable complexes, and MD simulations (four iterative 50 ns and 100 ns) demonstrated stable RMSD and RMSF profiles, persistent key interactions, and consistent ligand compactness for all three candidates. Elacestrant showed the most favorable binding free energy (ΔG_bind_ = −67.98 kcal/mol) and the greatest dynamic stability. Importantly, because Ct is an intracellular pathogen, we incorporated predicted cell-membrane permeability (QPPMDCK) into the screening: M Roflumilast and Tecovirimat showed excellent permeability, and Elacestrant exhibited moderate permeability, indicating that these candidates may reach the intracellular concentrations required to engage the CdsN interface. Consistent with their established clinical use, all three exhibit favorable pharmacokinetic properties and well-characterized safety profiles [87,89,94]. By contrast, none of the IMPPAT phytochemicals passed ADMET filtering despite favorable docking scores, most failing on predicted membrane permeability and intestinal absorption, reinforcing that binding affinity alone is insufficient for selecting drug-like antivirulence candidates.

This study is nonetheless constrained by its exclusively computational nature. The Ct T3SS model is built from homology- and template-based predictions rather than experimentally resolved structures, and the screening relies on docking, MM-GBSA, and MD simulations, which cannot fully capture in vivo efficacy, target selectivity, intracellular bioavailability, or host-cell toxicity. Accordingly, experimental validation will be essential. Future work should resolve the Ct T3SS subcomplex structures by cryo-electron microscopy or crystallography, confirm disruption of the CdsN oligomerization interface through ATPase-activity and oligomerization assays, and assess antivirulence efficacy in cell-based models. Collectively, our work provides the first molecular-level architecture of the Ct T3SS and a rational basis for targeting protein-protein interfaces in bacterial secretion systems, advancing the development of antivirulence therapeutics against chlamydial infection.

## 5. Conclusion

In this study, we present the first molecular-level model of the complete Ct T3SS, assembling all 13 constituent proteins into five subcomplexes that exhibit a highly conserved architecture despite low sequence identity with their Gram-negative homologs. While the predicted subcomplexes showed high structural and energetic reliability, their confirmation will require experimental structure determination. Exploiting this model, structure-based virtual screening against the CdsN ATPase oligomerization interface identified three FDA-approved drugs-M Roflumilast, Elacestrant, and Tecovirimat-that formed stable, high-affinity complexes with two adjacent CdsN monomers, with Elacestrant showing the most favorable binding free energy and dynamic stability. Although these findings are based solely on in silico analyses and warrant experimental validation, our work provides a rational framework for targeting PPI interfaces in bacterial secretion systems and advances the development of antivirulence strategies against chlamydial infection.

## Acknowledgements

AP is a recipient of senior research fellowship from the Department of Biotechnology (Grant number: DBT/2023-24/UOD/2326), Government of India. JK is a recipient of junior research fellowship from the University Grants Commission (Reference number: 231620067077), Government of India.

## Author contributions statement

**Amisha Panda:** methodology, software, investigation, data analysis, data curation, writing—original draft preparation. **Jahnvi Kapoor:** data analysis, visualization, writing—reviewing and editing. **Raman Rajagopal:** methodology, writing—reviewing and editing. **Sanjiv Kumar:** conceptualization, methodology, software, writing—reviewing and editing, supervision. **Anannya Bandyopadhyay:** methodology, data analysis, writing—original draft preparation, writing—reviewing and editing, supervision. All authors reviewed the final draft of the manuscript. All authors contributed to the manuscript revision, read and approved the submitted version.

## Conflict of interest

The authors declare no competing financial interest.

## Funding

This research was funded by Institute of Eminence (IOE), University of Delhi grant (Ref. No./IoE/2024-25/12/FRP) to RR and AB.

## Data Availability Statement

The atomic coordinate files (*.pdb) supporting the findings of this study have been deposited in the Zenodo public repository, accessible at https://doi.org/10.5281/zenodo.20741936. The Biopython-based script has been deposited in the Zenodo public repository, accessible at https://zenodo.org/records/18679384.

## Notes

### Competing Interest Statement

The authors have declared no competing interest.

### Summary of Updates

Title and Abstract revised; Introduction revised; Methods updated with additional 50 ns simulations; Results and Discussion revised accordingly; Figures 1, 6 and 8 revised; Tables updated; Supplemental file updated.

