## Supplemental material for "First Molecular-Level Model of the *Chlamydia trachomatis* Injectisome and Targeting CdsN ATPase Oligomerization for Antivirulence Therapy"

**Supplementary Tables:**

**Table S1:** Cross-validation of CdsC, CdsD, and CdsF models generated using five different modeling tools.

| **Structure Prediction Tools** | **Confidence scores of CdsC models** | **Confidence scores of CdsD models** | **Confidence scores of CdsF models** |
| --- | --- | --- | --- |
| **AlphaFold 3** | 0.42 | 0.37 | 0.49 |
| **ESMFold** | 0.410 | 0.331 | 0.752 |
| **RoseTTAFold2** | 0.524 | 0.606 | 0.472 |
| **SWISS-MODEL** | 0.66 | 0.65 | 0.82 |
| **TrRosetta** | 0.477 | 0.441 | 0.817 |
| **Structural alignment using US-Align** | | | |
| **RMSD (Å)** | 5.40 | 5.82 | 1.92 |

**Table S2:** Ramachandran plot analysis of the modeled Ct T3SS subcomplexes using Ramplot.

| **T3SS constituent proteins** | **Residues in allowed regions (%)** | **Residues in disallowed regions (%)** |
| --- | --- | --- |
| **Needle filament** |  |  |
| CdsF | 99.57 % | 0.44 % |
| **Basal body** |  |  |
| CdsC | 97.33 % | 2.67 % |
| CdsD-J | 99.29 % | 0.72 % |
| **Export apparatus** |  |  |
| CdsRST | 99.12 % | 0.87 % |
| CdsU | 100 % | 0 % |
| CdsV | 97.7 % | 2.30 % |
| **ATPase complex** |  |  |
| CdsN | 100 % | 0 % |
| CdsL | 100 % | 0 % |
| **Sorting platform** |  |  |
| CdsQ | 99.73 % | 0.27 % |

**Table S3:** Structure-based virtual screening: Docking scores and ADMET properties of eDrug3D compounds targeting the predicted hotspot residues of the CdsN oligomerization interface.

| **Drug ID** | **Name** | **Docking Score** | **SASA^a^** | **QPPMDCK^b^ (nm/s)** | **QPPCaco^c^**  **(nm/s)** | **QPlogPo/w^d^**  **(Lipophilicity)** | **donorHB** | **acceptHB** | **QPlogHERG^e^** | **Rule Of Five (Lipinski Rule)** | **Percentage of Human Oral Absorption** |
| --- | --- | --- | --- | --- | --- | --- | --- | --- | --- | --- | --- |
|  |  |  |  |  | **> 500** | **-2 to 6.5** | **≤ 5** | **≤ 10** | **< −5 = safer** | **≤ 1** | **> 80% = good** |
| 2040 | Difelikefalin | -9.97 | 1022.866 | 0.018 | 0.014 | -1.722 | 7.75 | 13.25 | -1.611 | 3 | 0 |
| 820 | Valrubicin | -8.881 | 992.449 | 32.367 | 11.147 | 2.23 | 3 | 16.6 | -4.79 | 2 | 32.824 |
| 774 | Ritonavir | -8.488 | 1179.223 | 264.313 | 103.441 | 6.757 | 3.25 | 10.95 | -5.797 | 3 | 63.69 |
| 969 | Gentamicin | -7.966 | 754.297 | 0.528 | 1.346 | -3.702 | 11 | 16.95 | -8.122 | 2 | 0 |
| 1863 | Abemaciclib | -7.947 | 898.096 | 131.581 | 98.633 | 4.507 | 1 | 9 | -8.227 | 1 | 76.065 |
| 739 | Acarbose | -7.929 | 802.791 | 0.055 | 0.199 | -6.628 | 14 | 32.1 | -4.929 | 3 | 0 |
| 1572 | Lomitapide | -7.804 | 1082.719 | 7268.587 | 503.85 | 8.93 | 2 | 7 | -7.951 | 2 | 100 |
| 1857 | Macimorelin | -7.784 | 746.74 | 17.627 | 16 | 1.948 | 4.25 | 5.75 | -2.631 | 1 | 46.947 |
| 2014 | Berotralstat | -7.729 | 942.453 | 37.981 | 11.442 | 4.944 | 4 | 7.5 | -9.123 | 1 | 61.878 |
| 2048 | M mobocertinib | -7.717 | 937.533 | 87.682 | 183.736 | 4.732 | 3 | 10.75 | -7.387 | 2 | 69.262 |
| 1898 | Plazomicin | -7.71 | 908.274 | 0.024 | 0.076 | -5.809 | 14 | 21.9 | -7.62 | 3 | 0 |
| 1076 | Paromomycin | -7.695 | 809.225 | 0.004 | 0.013 | -8.689 | 18 | 28.8 | -8.195 | 3 | 0 |
| 1023 | Netilmicin | -7.673 | 754.485 | 0.379 | 0.99 | -3.486 | 11 | 16 | -8.259 | 2 | 0 |
| 1898 | Plazomicin | -7.668 | 885.845 | 0.027 | 0.068 | -5.867 | 14 | 21.9 | -7.423 | 3 | 0 |
| 2046 | Mobocertinib | -7.611 | 965.218 | 181.073 | 359.393 | 5.341 | 2 | 10.75 | -7.48 | 3 | 65.085 |
| 968 | Neomycin | -7.609 | 829.215 | 0.001 | 0.003 | -9.487 | 19 | 28.1 | -8.988 | 3 | 0 |
| 1898 | Plazomicin | -7.601 | 924.044 | 0.034 | 0.077 | -5.7 | 14 | 21.9 | -7.807 | 3 | 0 |
| 602 | Clofazimine | -7.598 | 799.48 | 10000 | 5866.52 | 7.972 | 1 | 3 | -7.521 | 1 | 100 |
| 249 | Phenylpropanolamine | -7.58 | 373.994 | 214.304 | 420.021 | 0.597 | 3 | 2.7 | -4.534 | 0 | 77.394 |
| 1013 | Amikacin | -7.521 | 821.695 | 0.001 | 0.003 | -8.412 | 17 | 26.9 | -6.109 | 3 | 0 |
| 809 | Montelukast | -7.505 | 889.055 | 388.884 | 214.969 | 8.922 | 2 | 4.25 | -5.078 | 2 | 95.012 |
| 1863 | Abemaciclib | -7.488 | 879.304 | 121.924 | 90.92 | 4.355 | 1 | 9 | -8.014 | 1 | 74.54 |
| 2151 | Lazertinib | -7.478 | 934.606 | 199.662 | 393.401 | 4.502 | 2 | 11.45 | -8.157 | 2 | 73.83 |
| 1271 | Indinavir | -7.421 | 881.875 | 19.257 | 31.172 | 2.018 | 4 | 13.9 | -4.822 | 1 | 52.539 |
| 2047 | M mobocertinib | -7.411 | 1007.413 | 91.221 | 190.586 | 5.306 | 3 | 10.25 | -7.881 | 3 | 59.95 |
| 1238 | Bitolterol | -7.348 | 809.301 | 128.715 | 262.082 | 4.55 | 2 | 7.7 | -7.253 | 0 | 96.871 |
| 1314 | Tobramycin | -7.269 | 704.895 | 0.009 | 0.031 | -6.84 | 15 | 20.3 | -7.965 | 2 | 0 |
| 1074 | Streptomycin | -7.214 | 783.071 | 0.022 | 0.084 | -5.709 | 16 | 25.25 | -5.016 | 3 | 0 |
| 1787 | Isavuconazonium | -7.213 | - | **-** | **-** | **-** | **-** | **-** | **-** | **-** | **-** |
| 1881 | Tezacaftor | -7.187 | 777.143 | 913.705 | 409.435 | 3.649 | 4 | 9.6 | -5.725 | 1 | 82.105 |
| **1537** | **M roflumilast** | **-7.165** | **648.514** | **8784.547** | **1605.376** | **5.004** | **1** | **4.25** | **-5.424** | **1** | **100** |
| 1524 | Ticagrelor | -7.132 | 801.681 | 176.742 | 96.797 | 2.728 | 4 | 11.3 | -5.4 | 1 | 65.505 |
| **2081** | **Elacestrant** | **-7.118** | **776.187** | **361.172** | **680.758** | **6.391** | **2** | **4** | **-6.229** | **1** | **100** |
| 1861 | M netarsudil | -7.116 | 617.288 | 30.773 | 69.741 | 2.052 | 3 | 5.7 | -6.901 | 0 | 71.954 |
| 1318 | Kanamycin | -7.108 | 672.028 | 0.009 | 0.031 | -6.826 | 15 | 22.7 | -6.802 | 2 | 0 |
| 2070 | Adagrasib | -7.106 | 939.834 | 232.837 | 194.344 | 5.286 | 0 | 10 | -7.508 | 2 | 72.938 |
| 1391 | Glutathione disulfide | -7.104 | 901.529 | 0 | 0 | -5.49 | 9 | 18 | 5.852 | 3 | 0 |
| 1787 | Isavuconazonium | -7.077 | - | **-** | **-** | **-** | **-** | **-** | **-** | **-** | **-** |
| 1591 | M enzalutamide | -7.058 | 679.355 | 593.804 | 120.824 | 3.46 | 2 | 7 | -5.337 | 0 | 84.472 |
| 1843 | Telotristat ethyl | -7.029 | 851.209 | 165.892 | 44.582 | 5.421 | 4 | 6.5 | -6.753 | 2 | 62.287 |

**^#^**Values shown in red indicate the respective threshold limits for each pharmacokinetic property.

^a^SASA (solvent-accessible surface area) describes the extent of a molecular surface exposed to the solvent.

^b^QPPMDCK (nm/s) predicts cellular membrane permeability and estimates the ability of drug molecules to penetrate cell membrane.

^c^QPPCaco (nm/s) defines the predicted Caco-2 cell permeability, which indicates intestinal absorption potential.

^d^QPlogPo/w represents the predicted water partition coefficient and reflects the compound lipophilicity.

^e^QPlogHERG is the predicted potential for hERG channel inhibition, which is associated with cardiac toxicity risk.

**Table S4:** Structure-based virtual screening: Docking scores and ADMET properties of eDrug3D compounds targeting the druggable pocket predicted within the CdsN oligomerization interface.

| **Drug ID** | **Name** | **Docking Score** | **SASA^a^** | **QPPMDCK^b^ (nm/s)** | **QPPCaco^c^ (nm/s)** | **QPlogPo/w^d^ (Lipophilicity)** | **donorHB** | **acceptHB** | **QPlogHERG** | **Rule Of Five (Lipinski Rule)** | **Percentage of Human Oral Absorption** |
| --- | --- | --- | --- | --- | --- | --- | --- | --- | --- | --- | --- |
|  |  |  |  |  | **> 500 = good** | **-2 to 6.5** | **≤ 5** | **≤ 10** | **> −5 = safer** | **≤ 1** | **> 80% = good** |
| 1286 | Frovatriptan | -8.826 | 496.783 | 39.877 | 88.636 | 0.853 | 4 | 4 | -5.09 | 0 | 66.798 |
| 2040 | Difelikefalin | -8.476 | 1172.005 | 0.015 | 0.009 | -0.932 | 7.75 | 13.25 | -3.062 | 3 | 0 |
| 116 | Metharbital | -8.441 | 383.796 | 403.077 | 827.361 | 0.86 | 1 | 4 | -2.359 | 0 | 84.205 |
| 1644 | Adefovir | -8.232 | 504.265 | 0.685 | 1.4 | -0.657 | 4 | 10.7 | -1.048 | 0 | 25.718 |
| 359 | Allopurinol | -7.993 | 298.28 | 76.192 | 177.156 | -0.569 | 1 | 4 | -2.973 | 0 | 63.853 |
| 1393 | Inamrinone | -7.878 | 393.424 | 103.956 | 236.154 | 0.176 | 2.5 | 5 | -4.185 | 0 | 70.45 |
| 1128 | Ethosuximide | -7.814 | 316.389 | 493.599 | 997.923 | 0.711 | 1 | 3 | -1.956 | 0 | 84.785 |
| 582 | Tranexamic acid | -7.657 | 371.789 | 10.834 | 21.252 | -1.287 | 3 | 3 | -1.736 | 0 | 43.168 |
| 232 | Chlorzoxazone | -7.597 | 323.37 | 1195.239 | 981.751 | 1.092 | 1 | 3 | -3.195 | 0 | 86.892 |
| 76 | Aminosalicylic acid | -7.430 | 332.733 | 13.301 | 28.211 | 0.354 | 2.5 | 2.75 | -1.504 | 0 | 54.979 |
| 1027 | Cefoperazone | -7.410 | 978.39 | 0.151 | 0.101 | 0.625 | 2.25 | 15.5 | -1.908 | 2 | 0 |
| 1664 | M tazarotene | -7.364 | 613.157 | 125.184 | 150.712 | 4.463 | 1 | 3.5 | -3.864 | 0 | 92.063 |
| 2040 | Difelikefalin | -7.363 | 1145.456 | 0.018 | 0.014 | -1 | 7.75 | 13.25 | -2.804 | 3 | 0 |
| 1436 | Lanreotide | -7.289 | 1457.983 | 0.01 | 0.001 | 0.468 | 9.75 | 19.2 | 0.756 | 3 | 0 |
| 1399 | Mesalamine | -7.207 | 332.983 | 13.299 | 28.207 | 0.356 | 2.5 | 2.75 | -1.511 | 0 | 54.986 |
| 754 | Lamivudine | -7.143 | 416.032 | 164.467 | 215.053 | -0.479 | 3 | 7.9 | -3.498 | 0 | 65.887 |
| **1889** | **Tecovirimat** | **-7.127** | **605.725** | **1812.978** | **849.205** | **3.9** | **0.25** | **4.75** | **-5.167** | **0** | **100** |
| 35 | Homatropine methylbromide | -7.111 | - | **-** | **-** | **-** | **-** | **-** | **-** | **-** | **-** |
| 88 | Methimazole | -7.106 | 292.791 | 5629.055 | 4036.873 | 0.727 | 1 | 2.5 | -2.953 | 0 | 95.747 |
| 1755 | M allopurinol | -7.065 | 286.964 | 26.431 | 66.523 | -1.842 | 3 | 4.5 | -2.215 | 0 | 48.786 |
| 177 | Hydroxyamphetamine | -7.036 | 375.573 | 108.846 | 224.424 | 0.663 | 3 | 1.75 | -4.378 | 0 | 72.905 |
| 9 | Theophylline | -7.005 | 386.501 | 133.19 | 297.006 | -0.02 | 1 | 5 | -3.594 | 0 | 71.089 |
| 149 | Oxtriphylline | -7.005 | 386.501 | 133.19 | 297.006 | -0.02 | 1 | 5 | -3.594 | 0 | 71.089 |

**^#^**Values shown in red indicate the respective threshold limits for each pharmacokinetic property.

^a^SASA (solvent-accessible surface area) describes the extent of a molecular surface exposed to the solvent.

^b^QPPMDCK (nm/s) predicts cellular membrane permeability and estimates the ability of drug molecules to penetrate cell membrane.

^c^QPPCaco (nm/s) defines the predicted Caco-2 cell permeability, which indicates intestinal absorption potential.

^d^QPlogPo/w represents the predicted water partition coefficient and reflects the compound lipophilicity.

^e^QPlogHERG is the predicted potential for hERG channel inhibition, which is associated with cardiac toxicity risk.

**Table S5:** Structure-based virtual screening: Docking scores and ADMET properties of IMPPAT compounds targeting the predicted hotspot residues of the CdsN dimer.

| **Phytochemical Identifier** | **Phytochemical Name** | **Docking score** | **SASA^a^** | **QPPMDCK^b^**  **(nm/s)** | **QPPCaco^c^ (nm/s)** | **QPlogPo/w^d^ (Lipophilicity)** | **donorHB** | **acceptHB** | **QPlogHERG^e^** | **Rule Of Five (Lipinski Rule)** | **Percentage of Human Oral Absorption** |
| --- | --- | --- | --- | --- | --- | --- | --- | --- | --- | --- | --- |
|  |  |  |  | **<25 poor;**  **25-500 moderate;**  **>500 excellent** | **> 500** | **-2 to 6.5** | **≤ 5** | **≤ 10** | **< −5 = safer** | **≤ 1** | **> 80% = good** |
| IMPHY012839 | Lupin isoflavone D | -7.458 | 611.926 | 55.402 | 131.923 | 2.477 | 3 | 5.25 | -5.284 | 0 | 79.4 |
| IMPHY002401 | Holarricine | -7.006 | 576.443 | 3.279 | 8.002 | 0.637 | 4 | 6.7 | -4.847 | 0 | 46.839 |

**^#^**Values shown in red indicate the respective threshold limits for each pharmacokinetic property.

^a^SASA (solvent-accessible surface area) describes the extent of a molecular surface exposed to the solvent.

^b^QPPMDCK (nm/s) predicts cellular membrane permeability and estimates the ability of drug molecules to penetrate cell membrane.

^c^QPPCaco (nm/s) defines the predicted Caco-2 cell permeability, which indicates intestinal absorption potential.

^d^QPlogPo/w represents the predicted water partition coefficient and reflects the compound lipophilicity.

^e^QPlogHERG is the predicted potential for hERG channel inhibition, which is associated with cardiac toxicity risk.

**Table S6:** Structure-based virtual screening: Docking scores and ADMET properties of IMPPAT compounds targeting the druggable pocket predicted within the CdsN dimer.

| **Phytochemical Identifier** | **Phytochemical Name** | **Docking score** | **SASA^a^** | **QPPMDCK^b^**  **(nm/s)** | **QPPCaco^c^**  **(nm/s)** | **QPlogPo/w^d^**  **(Lipophilicity)** | **donorHB** | **acceptHB** | **QPlogHERG^e^** | **Rule Of Five (Lipinski Rule)** | **Percentage of Human Oral Absorption** |
| --- | --- | --- | --- | --- | --- | --- | --- | --- | --- | --- | --- |
|  |  |  |  | **<25 poor;**  **25-500 moderate;**  **>500 excellent** | **> 500** | **-2 to 6.5** | **≤ 5** | **≤ 10** | **< −5 = safer** | **≤ 1** | **> 80% = good** |
| IMPHY005259 | 7-Hydroxy-2-oxindole-3-acetic acid | -8.048 | 392.818 | 10.921 | 23.508 | 0.217 | 3 | 5.25 | -1.775 | 0 | 52.757 |
| IMPHY000145 | (+)-Isofebrifugine | -7.998 | 549.462 | 45.332 | 99.799 | 0.67 | 1 | 8.2 | -5.533 | 0 | 66.647 |
| IMPHY011989 | (-)- Afzelechin | -7.758 | 502.507 | 67.772 | 158.965 | 1.154 | 4 | 4.7 | -4.916 | 0 | 73.1 |
| IMPHY012224 | (-)-Catechin | -7.735 | 515.615 | 22.7 | 57.785 | 0.492 | 5 | 5.45 | -4.879 | 0 | 61.359 |
| IMPHY006061 | Homogentisic acid | -7.326 | 358.697 | 17.072 | 35.54 | 0.406 | 3 | 3.5 | -1.669 | 0 | 57.076 |
| IMPHY000891 | Miraxanthin iiis | -7.239 | 630.433 | 0.816 | 1.707 | 1.977 | 4 | 7.25 | -2.155 | 0 | 42.675 |
| IMPHY010119 | (-)-Leucofisetinidin | -7.186 | 513.935 | 25.755 | 64.947 | 0.283 | 5 | 6.4 | -4.968 | 0 | 61.042 |

**^#^**Values shown in red indicate the respective threshold limits for each pharmacokinetic property.

^a^SASA (solvent-accessible surface area) describes the extent of a molecular surface exposed to the solvent.

^b^QPPMDCK (nm/s) predicts cellular membrane permeability and estimates the ability of drug molecules to penetrate cell membrane.

^c^QPPCaco (nm/s) defines the predicted Caco-2 cell permeability, which indicates intestinal absorption potential.

^d^QPlogPo/w represents the predicted water partition coefficient and reflects the compound lipophilicity.

^e^QPlogHERG is the predicted potential for hERG channel inhibition, which is associated with cardiac toxicity risk.

**Table S7:** Prime MM/GBSA-derived binding free energy (ΔG) parameters of selected ligands docked against the CdsN dimer.

| **Drug ID** | **Drug name** | **ΔG_cou_** | **ΔG_cov_** | **ΔG_lipo_** | **ΔG_vdW_** |
| --- | --- | --- | --- | --- | --- |
| 1537 | M Roflumilast | 6.46 | 1.06 | -15.42 | -41.3 |
| 2081 | Elacestrant | -16.34 | 21.19 | -32.98 | -94.27 |
| 1889 | Tecovirimat | -42.43 | 12.4 | -12.88 | -39.8 |

ΔG_cou_, Coulombic energy (kcal/mol); ΔG_cov_, covalent energy (kcal/mol); ΔG_lipo_, hydrophobic energy (kcal/mol); ΔG_vdW_, van der Waals energy (kcal/mol).

**Supplementary Figures:**

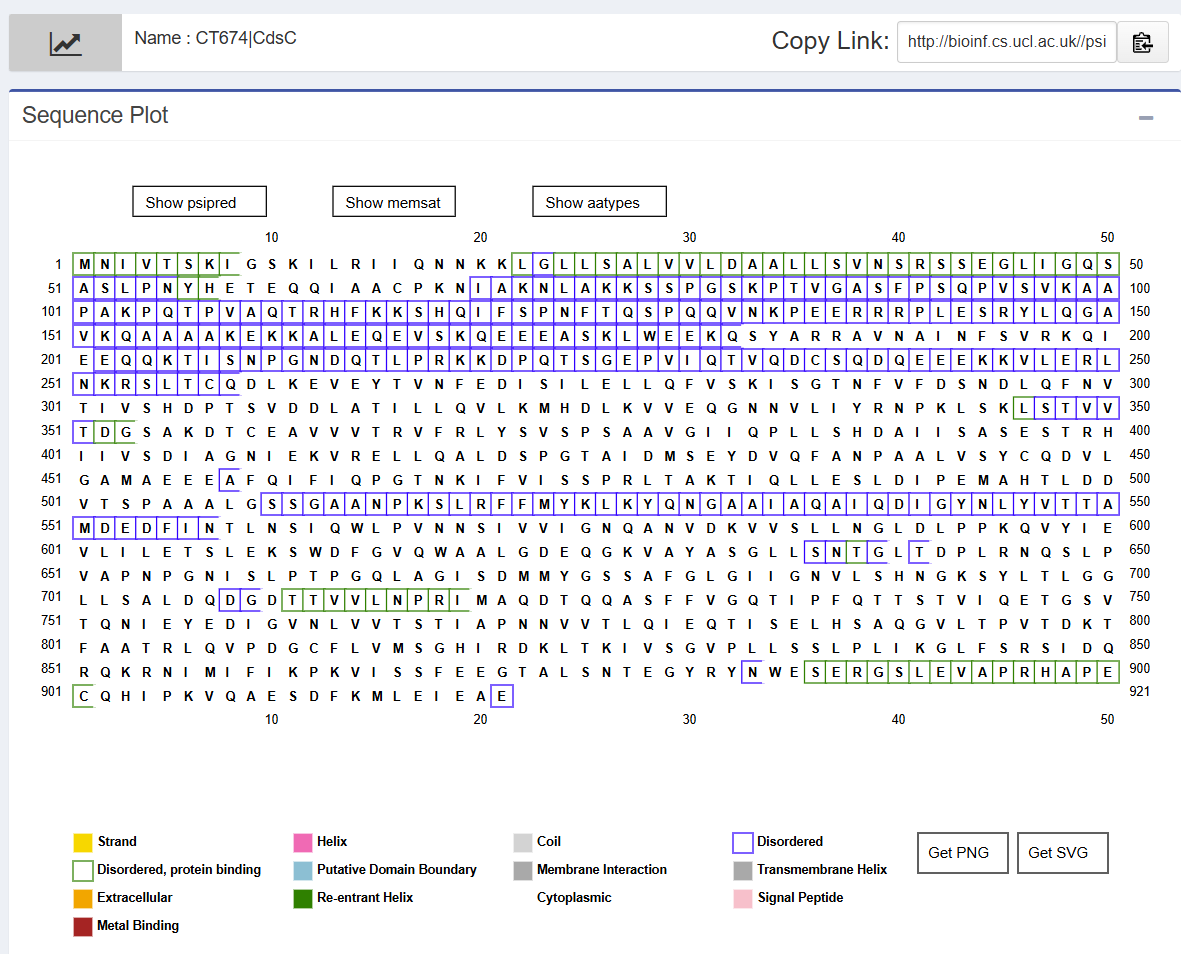

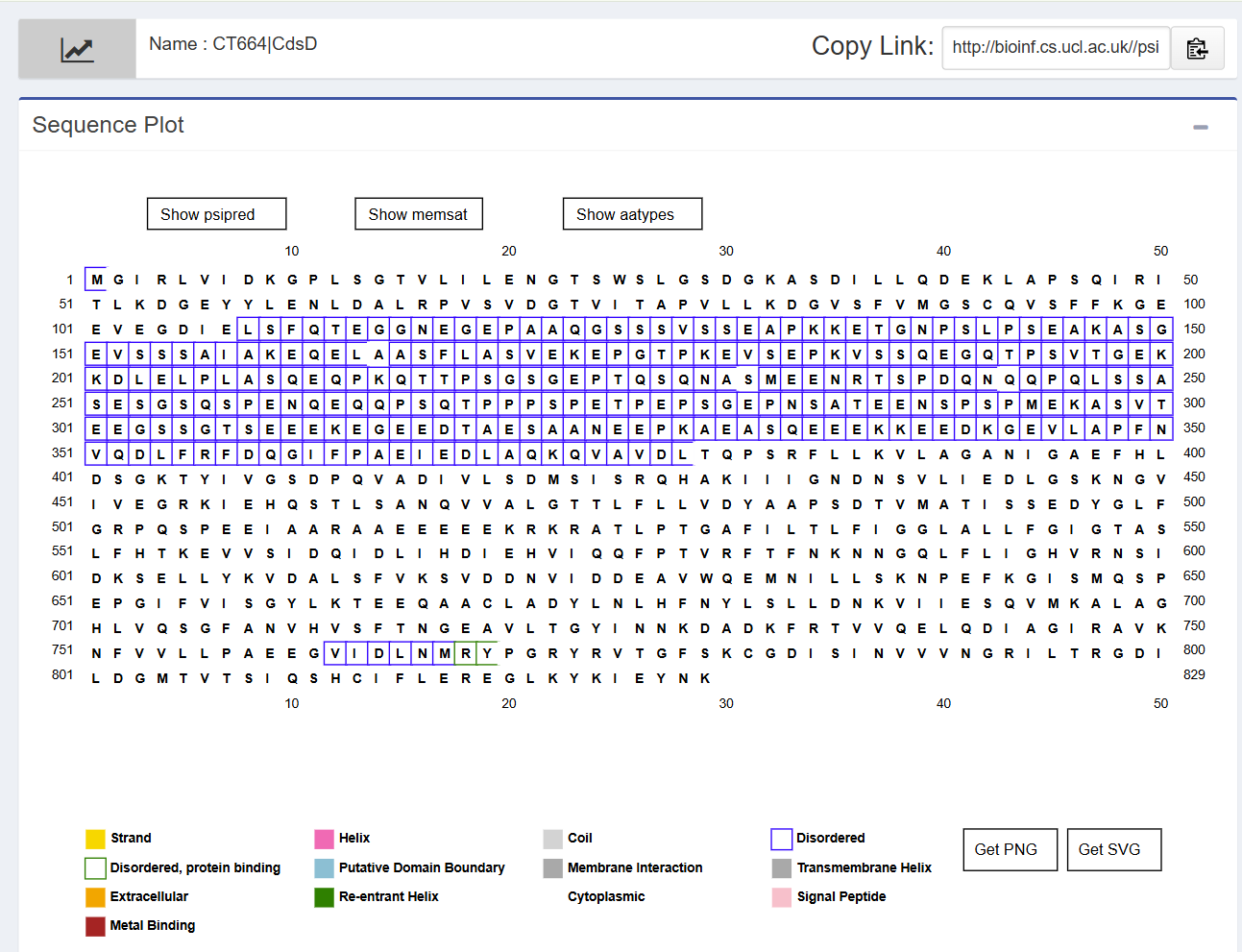

**Figure S1:** **Intrinsically disordered regions predicted for the proteins. (a)** CdsC and **(b)** CdsD

**(b)**

**(a)**

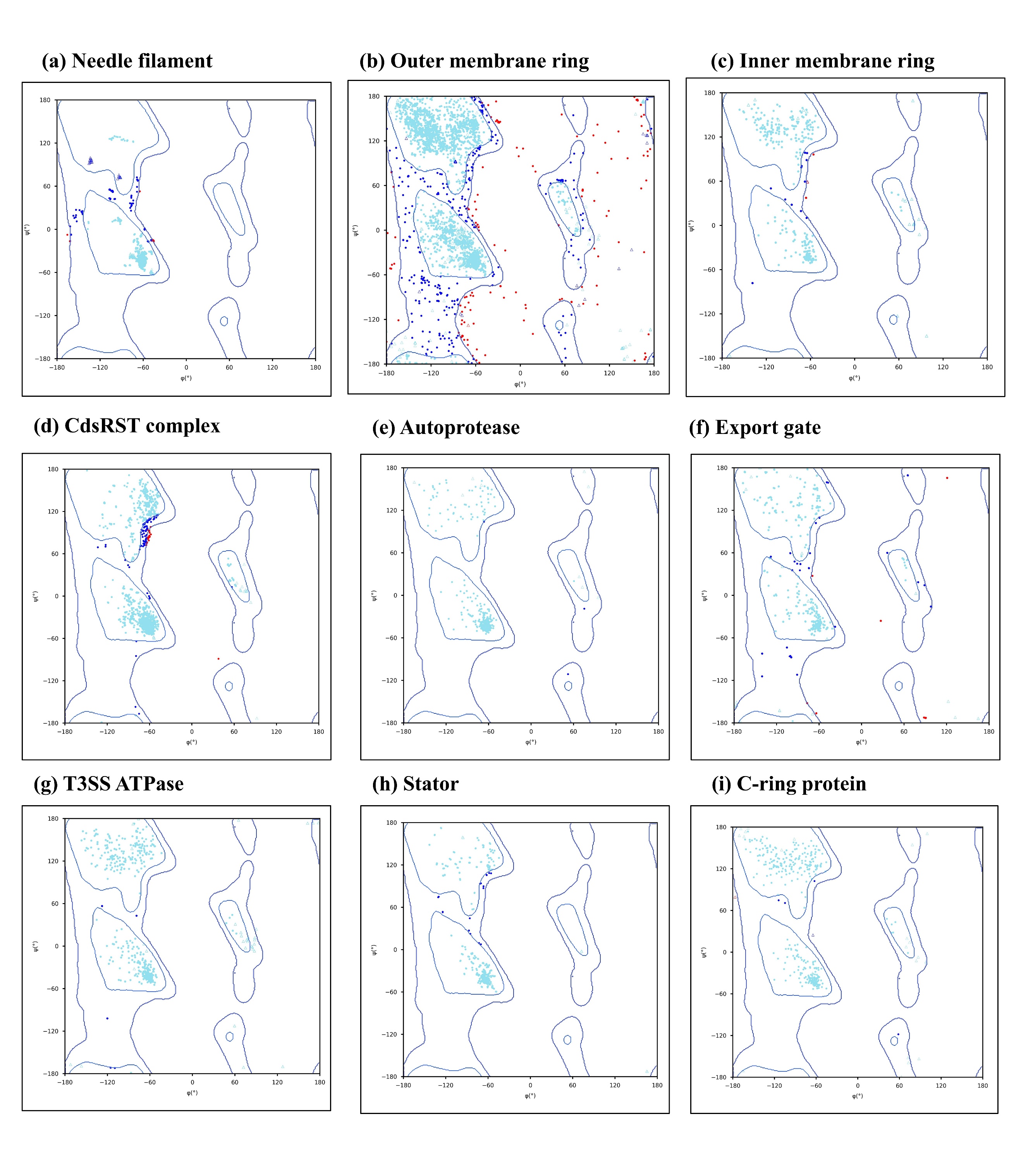

**Figure S2: Ramachandran plot analysis of Ct T3SS constituent proteins using Ramplot.** Ramachandran plot represents the backbone dihedral angles (φ and ψ) of amino acid residues for modeled Ct T3SS protein constituents. Green, blue, and red symbols correspond to favored, allowed, and disallowed regions, respectively, whereas dots represent non-glycine, and triangles represent glycine residues. The plots show the percentage of residues falling in favored regions for all the modeled T3SS components: **(a)** Needle filament, **(b)** Outer membrane ring, **(c)** Inner membrane ring, **(d)** CdsRST complex, **(e)** Autoprotease, **(f)** Export gate, **(g)** T3SS ATPase, **(h)** Stator, and **(i)** C-ring protein.

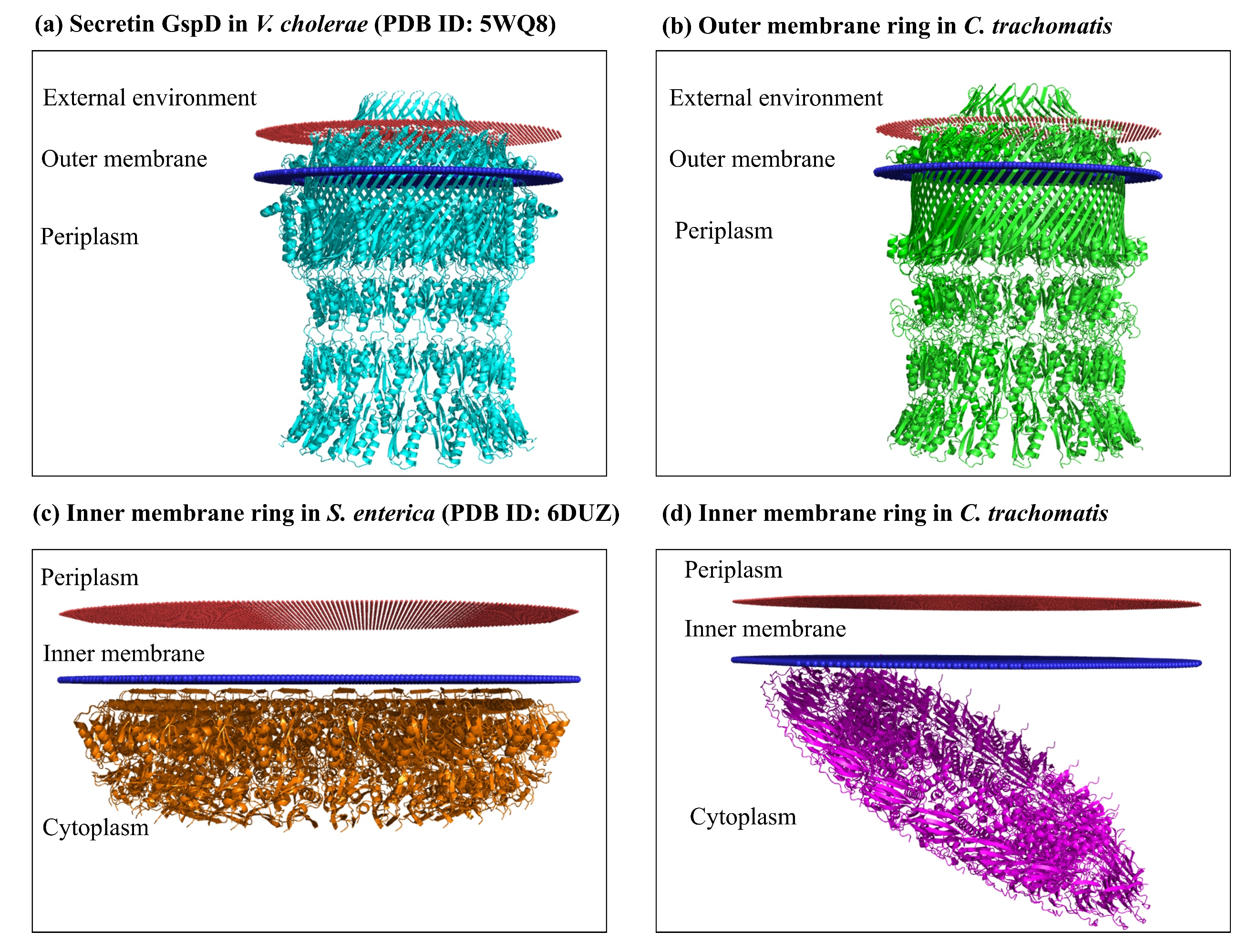

**Figure S3: Orientation of Ct T3SS subcomplexes and their corresponding homolog within the membrane predicted by the PPM server.** The membrane insertion and orientation of the outer membrane ring and inner membrane ring were predicted using the PPM server. **(a, b)** The outer membrane pore/ring of *V. cholerae* and Ct show similar overall topology, spanning the outer membrane (OM) with C-terminal domains forming a barrel in the OM and N-terminal domains extending towards the periplasmic side. **(c, d)** Inner membrane rings of *S. enterica* and Ct demonstrate somewhat more or less similar orientation patterns. In both the species, the protein is localized towards the cytoplasmic side.

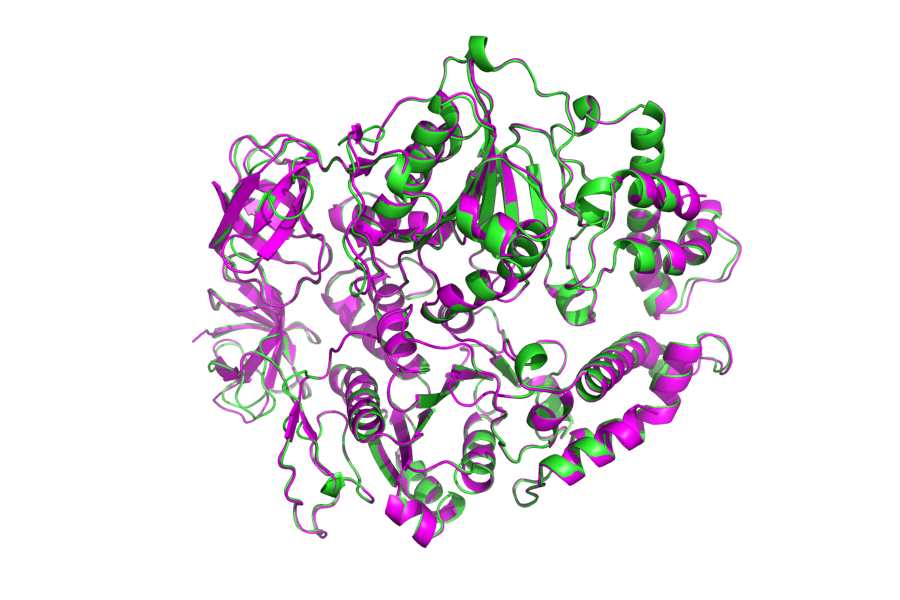

Ct CdsN dimer

*E. coli* EscN dimer

**Figure S4:** Structural alignment of the Ct CdsN dimer model with the *E. coli* EscN dimer crystal structure (PDB ID: 6NJP) using PyMOL.

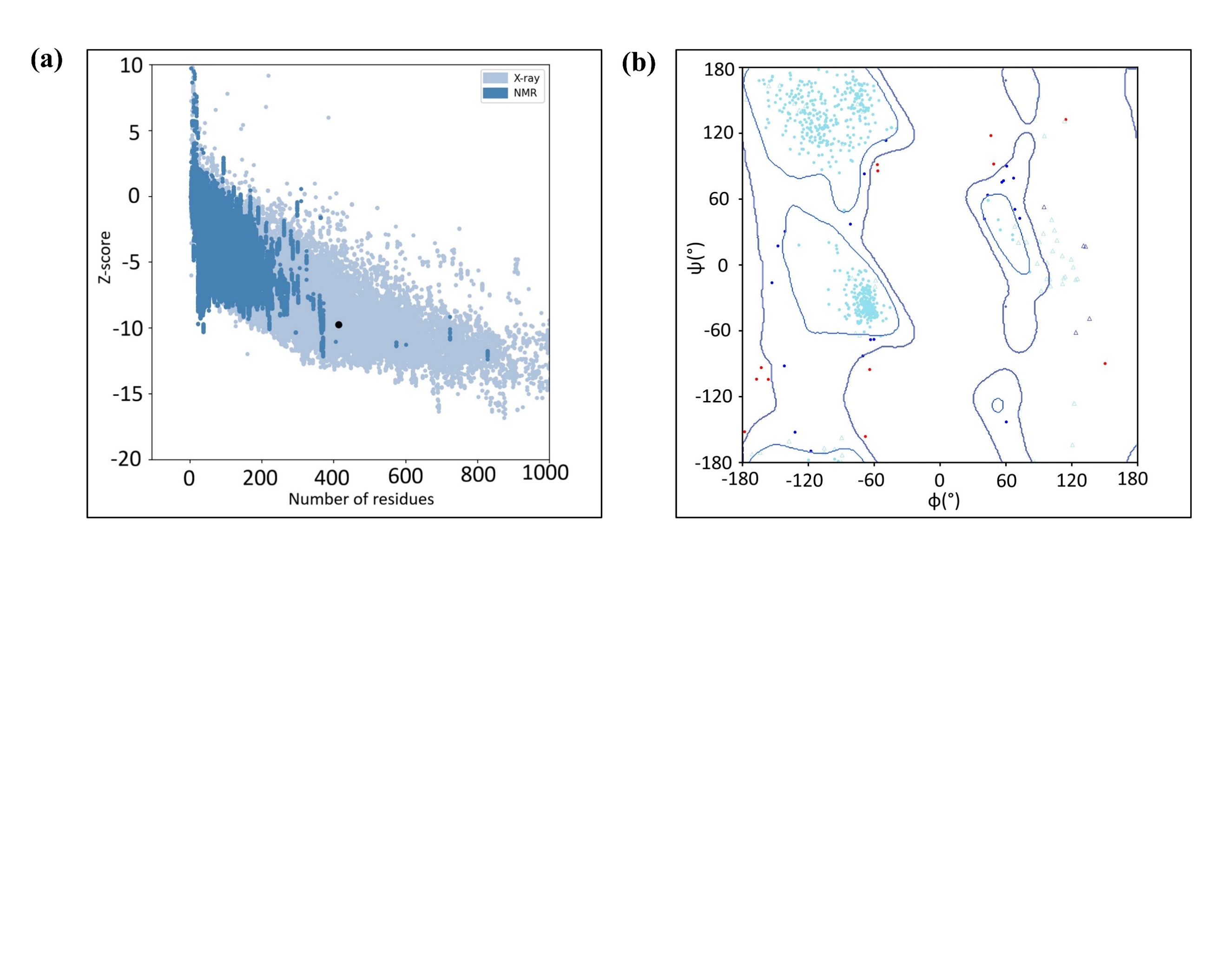

**Figure S5**: **Model quality assessment of Ct T3SS ATPase, CdsN dimer. (a)** Z-score plot generated using ProSA-web server, where the black dot corresponds to the query structure. The light blue region represents protein structures solved by X-ray crystallography, while the dark blue region corresponds to those determined by nuclear magnetic resonance (NMR) spectroscopy. **(b)** Ramachandran plot generated using the Ramplot webserver showing 95.355% of residues in the favored region. Green, blue, and red symbols correspond to favored, allowed, and disallowed regions, respectively; dots represent non-glycine, and triangles represent glycine residues.

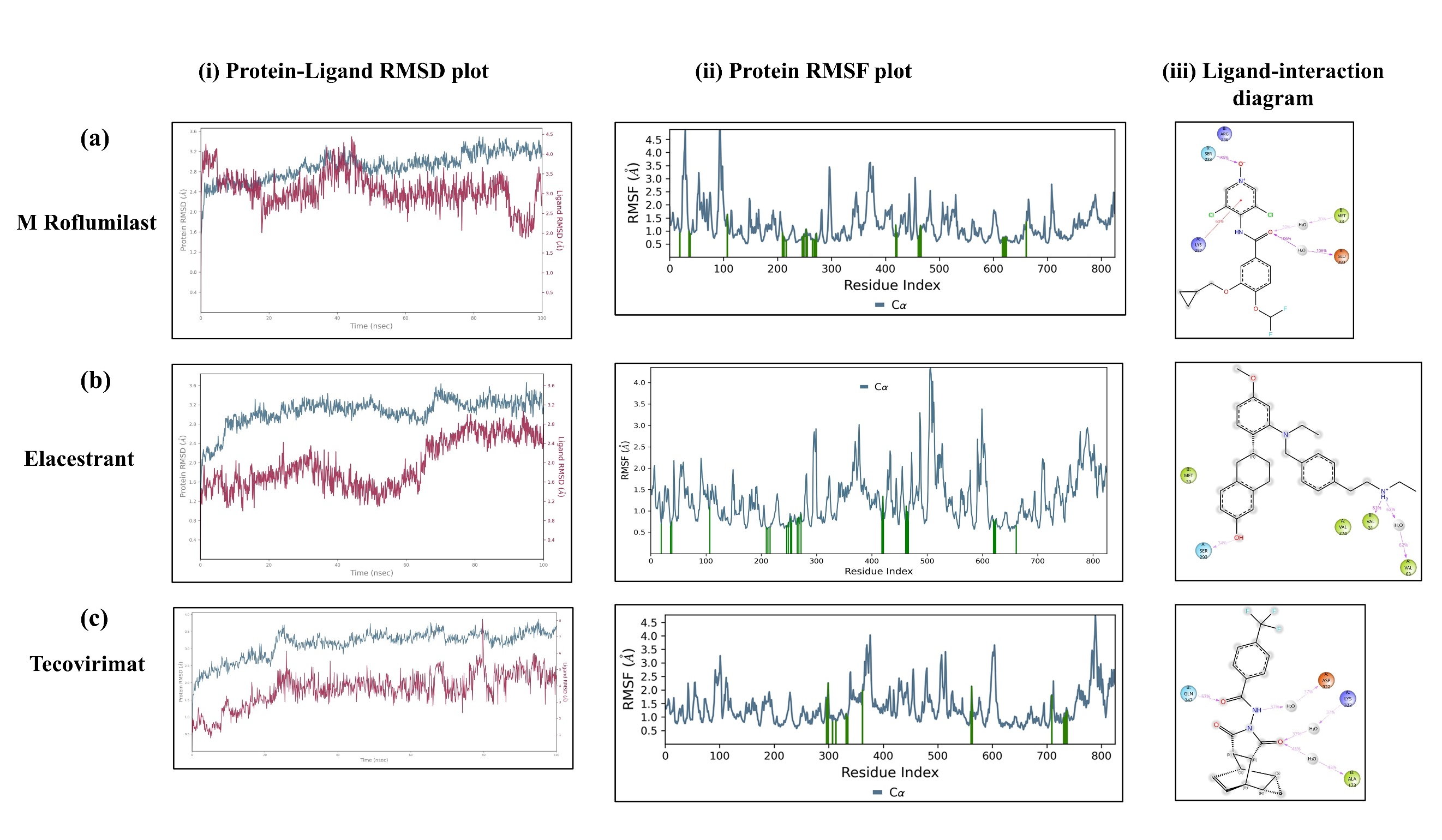

**Figure S6: Molecular dynamics simulation of the CdsN dimer-ligand complexes over 100 ns. (a-c)** CdsN dimer in complex with **(a)** M Roflumilast, **(b)** Elacestrant, and **(c)** Tecovirimat. For each complex: **(i)** Time-dependent protein-ligand root mean square deviation (RMSD) profile over 100 ns; **(ii)** Residue-wise root mean square fluctuation (RMSF) plot of the protein backbone (Cα atoms), where ligand-interacting residues are highlighted with green vertical bars; **(iii)** A two-dimensional schematic representation of detailed ligand atom interactions with the protein residues. Only interactions persisting for more than 30% of the simulation time (0-100 ns) are shown. Solid magenta arrows show hydrogen bonds, while dashed magenta arrows represent side-chain hydrogen bonds.

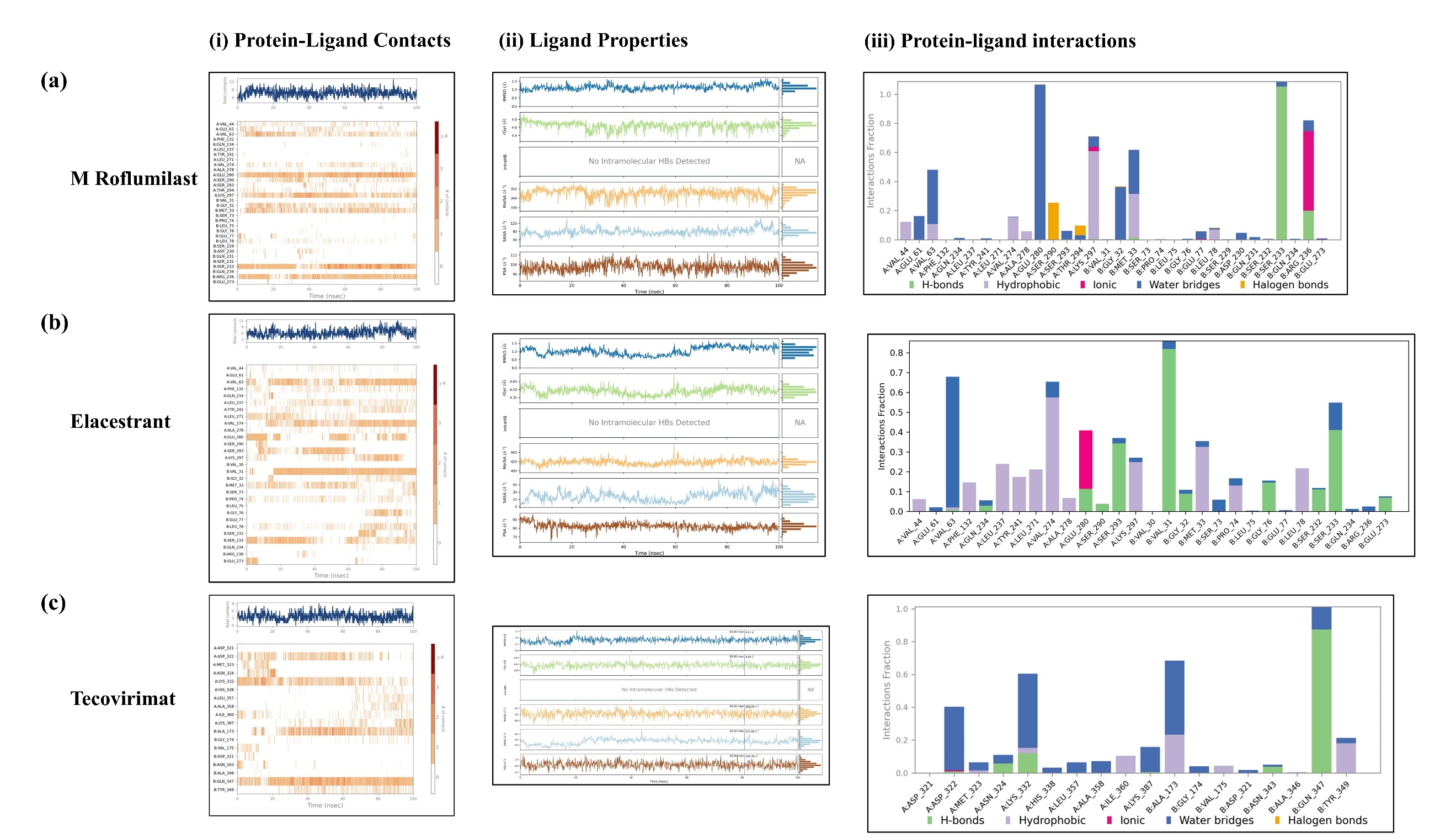
**Figure S7: Molecular dynamics simulation of the CdsN dimer-ligand complexes over 100 ns. (a-c)** CdsN dimer in complex with **(a)** M Roflumilast, **(b)** Elacestrant, and **(c)** Tecovirimat. For each complex: **(i)** A timeline representation of protein-ligand interactions (hydrogen bonds, hydrophobic contacts, ionic interactions, water bridges, and halogen bonds) is shown. **(ii)** Dynamics of ligand’s physicochemical properties during the simulation period. **(iii)** Protein-ligand interaction profile throughout the simulation, categorized into hydrogen bonds, hydrophobic contacts, ionic interactions, water bridges, and halogen bonds.
